# Catecholamine Precursor Modulation of Human Exploration: Evidence From a Large Gender-Balanced Sample

**DOI:** 10.1101/2025.03.18.643866

**Authors:** A. Brands, K. Knauth, D. Mathar, T. Rödder, K. Lisner, J. Peters

## Abstract

The catecholamine precursor tyrosine has been linked to improved cognitive performance, but investigations into decision-making and reinforcement learning processes known to be under catecholamine control are sparse. We examined the impact of a single dose of Tyrosine (2g) on reinforcement learning and exploration in a large (n=63) gender-balanced sample in a within-subjects preregistered study. Reinforcement learning performance was improved under Tyrosine, and computational modeling revealed that this performance increase was due to a stabilization of choice behavior reflected in increased value-driven exploitation. Further non-preregistered modeling analyses confirmed that accounting for higher-order perseveration substantially improved model fit, and substantiated the observation of increased value-driven exploitation under Tyrosine. Furthermore, it revealed a more fine-grained computational impact of Tyrosine, showing attenuated effects of directed exploration and value-independent perseveration. Supplementation with Tyrosine therefore improved reinforcement learning performance by stabilizing choice patterns in the service of optimizing reward accumulation. Results confirm that Tyrosine supplementation modulates specific computational mechanisms thought to be under catecholamine control.

## Introduction

Freely available supplements that promise enhancing effects on cognition are on the rise. Yet, if and how dietary supplements may affect cognitive function and performance is still poorly understood. Tyrosine (TYR) is the precursor of catecholamine neurotransmitters and is thought to facilitate the synthesis of dopamine, norepinephrine, and epinephrine (Bear et al., 2018). Cognitively demanding activities and stress increase catecholamine demand, which in turn leads to a decrease in TYR plasma availability (Jongkees et al., 2015; Kvetnansky et al., 2009; Lehnert et al., 1984). Thus, additional intake of TYR is assumed to provide the necessary resources to reach and maintain optimal performance under demanding conditions (Jongkees et al., 2015; Wurtman et al., 1980).

Such claims seem to be supported by findings on enhancing and protective effects of TYR on working memory (WM) and executive control (especially under conditions of heightened demand; Colzato et al., 2014; Hase et al., 2015; Jongkees et al., 2015). Nonetheless, despite the central role of the catecholamine neurotransmitter dopamine (DA) in reinforcement learning (RL) and decision-making (Namboodiri, 2024; Pessiglione et al., 2006; Taira & Sharpe, 2025; Westbrook et al., 2025; Wang et al., 2024)., comparatively little is known about potential TYR-related changes in these functions. An earlier report suggested that TYR may improve some aspects of decision-making (Mathar et al 2022), but this study was restricted to male participants, thus limiting generalizability. TYR effects on RL and decision-making are of considerable relevance, however, as both aberrant DA-functioning and RL deficits have been linked to several psychiatric disorders and maladaptive behaviors on the subclinical spectrum (Addicot et al., 2017; Chakroun et al., 2020; Cremer et al., 2023; Goschke 2014; Montague et al., 2012; Morris et al., 2016; Schwartenbeck et al., 2019; Wiehler et al., 2021). This resonates with the approach taken in the field of computational psychiatry, which investigates neurocomputational components underlying mental disorders (Voon et al., 2017; Huys et al., 2016; Montague et al., 2012; Wise et al., 2022; Yip et al., 2022). Especially the balance between more elaborate and cognitively costly *goal-directed* choice mechanisms on the one hand and simpler, less flexible *habitual* choice mechanisms, on the other, have emerged as highly relevant. Imbalances between these two broad systems are present across a range of disorders (Goschke 2014; Hauser et al., 2022; Hogeveen et al., 2022; Huys et al., 2016; 2021; Montague et al., 2012; Voon et al., 2017), and initial evidence points towards an impact of TYR (Mathar et al. 2022).

A central problem in RL is the tradeoff between exploration (sampling novel options for information gain) and exploitation (selecting known options for reward maximization) (Addicot et al., 2017; Cohen et al., 2007; Morris et al., 2016; Wilson et al., 2014). Dynamically changing environments require a balance of these strategies (Findling et al., 2021; Hogeveen et al., 2022; Wilson et al., 2014), since both overreliance on exploitation and excessive exploration can lead to performance decrements (Dubois & Hauser, 2021; Fox et al., 2020; Schulz & Gershman, 2019). Exploration can be further subdivided into (at least) two component processes: random vs. directed exploration (Gershman, 2018; 2019; Wilson et al., 2014, 2021). While random exploration is characterized by choice randomization, directed exploration is linked to a strategic exploration of uncertain options for information gain (Feng et al., 2021; Fox et al., 2020; Hogeveen et al., 2022, Wilson et al. 2021). This can be formalized by including an uncertainty-based “exploration bonus” in formal RL models (Schwartenbeck et al., 2019; Schulz & Gershman, 2019, Wiehler et al., 2021; Chakroun et al., 2020, Wilson et al,2021; Danwitz et al., 2024; Tuzsus et al., 2024).

Several mental disorders and subclinical variations thereof have been reliably associated with aberrant explore-exploit behavior (e.g. Addicot et al., 2017; Aylward et al., 2019; Dubois & Hauser, 2021; Gershman 2018; Goschke, 2014; Wiehler et al., 2021). Likewise, manipulations of the DA system have been linked to changes in exploration (Chakroun et al., 2020; Cremer et al., 2023; Domenech et al., 2020; Dubois et al., 2021; Kayser et al., 2015) and conceptually closely-related model-based control (Chakroun, et al., 2020; Mathar et al, 2022). However, the specific role of TYR in regulating exploration is currently unclear.

To address this issue, we preregistered a counterbalanced, placebo-controlled double-blind within-subject study to assess the impact of a single 2g dose of tyrosine (vs. placebo) in a gender-balanced sample. We opted for a comparatively large sample size (n=63 in total), considering mostly small samples sizes prevalent in the supplementation field. The preregistration and study information can be found at https://osf.io/4z39r/files/osfstorage.

## Methods

### Sample

The study procedure was approved by the ethics committee of the Faculty for Human Sciences of the University of Cologne. We pre-registered a sample size of 68 (34 subjects per group). Overall 69 subjects completed participation, four of which were excluded due to high BDI-scores and the fifth participant was excluded due to their acute self-reported health status (i.e. no sleep, high caffeine intake; see exclusion criteria as preregistered). We further excluded one more participant who showed nonsensical response patterns in one of the sessions. Thus, a total of N=63 (32 self-identified female) comprised the final sample. Participants had a mean age of M = 23.38 (SD = 5.30; range= 18 to 52 years). The vast majority (62 out of the final 63) had a degree in higher education and an average Body-Mass-Index (BMI; M = 22.93, SD = 3.60; range= 17.53 to 34.72). A descriptive summary of the self-reports is provided in the supplementary Table S1.

### Procedure

Participants participated in testing at the University of Cologne Biopsychology lab on two separate days. Sessions were spaced 3-5 days apart and took place in the same time slot each for every participant. Prior to participation all subjects gave their informed consent. On each testing day we first obtained baseline measures of physiological measures such as skin conductance response, heart rate and pupillometry. Individuals then consumed 200ml of orange juice containing either 2g of dissolved TYR (TheHut.com Ltd.) or PLC (microcrystalline cellulose), followed by a 60-minute waiting period in a separate room (with minimal external stimulating or distracting features; i.e. empty office room). Participants were instructed to abstain from drinking alcohol or consuming protein-rich food from the evening prior and up to each testing day in order to minimize external influences on TYR levels (Deijen, 2005).

Following the 60min waiting period, physiological measures were obtained again, and participants performed a short temporal discounting task (10-15min). Temporal discounting and physiological data were obtained following the procedures of Mathar et al. (2022) and were included as a direct replication, which will be reported elsewhere. This was followed by 300 trials of a four-armed restless bandit task (Chakroun et al., 2020, Wiehler et al., 2021, Daw et al., 2006) with changing option reward values over time (Figure 1). Values changed independently following Gaussian random walks. Participants could gain an additional payout of up to 4€ depending on task performance. Task and supplementation order were randomized and counterbalanced within each gender group and thus, also across the entire sample. At the end of the second testing day participants also completed a number of self-reports and sociodemographic questionnaires not relevant for the present report (see preregistration, Table S1).

**Figure 1.**
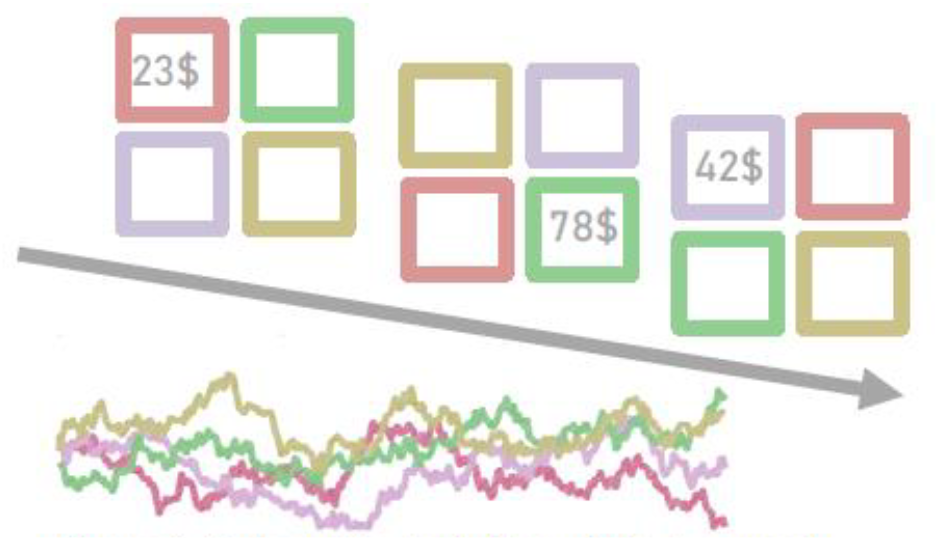
Schematic depiction of the 4-armed restless bandit task. On each trial, participants selected between four choice options marked by different colors. Following each choice, the reward associated with that option was displayed. Rewards varied independently according to Gaussian random walks (Wiehler et al., 2021; Daw et al., 2006; Chakroun et al., 2020) and were displayed in the form of hypothetical monetary amounts between 0 and 100€.

### Model-Agnostic Behavioral Data Analysis

As model-agnostic variables of interest, we examined median reaction times (RTs), the proportion of highest-value choices (%optimal), and participants’ tendency to switch between choice options (%switches). To this end, we set up linear mixed models with factors sex (male vs. female), condition (tyrosine vs. placebo) and their interaction as fixed effects. Subjects were included as random effects. This yielded three separate regression models for RT, %optimal, and %switches. Regression analyses using linear mixed models included sex and supplementation condition, along their interaction as fixed and subjects as random effects using the *lmerTest* package (Kuznetsova et al., 2017) in the statistical program R (R Development Team, 2020).

### Hierarchical Bayesian Modelling

In addition to the analysis of model-agnostic performance indices (see above) our main focus was on computational modelling using a hierarchical Bayesian approach (c.f. preregistered general procedure). In a first step, we compared a range of competing computational accounts of behavior, based on commonly-applied RL model variants (see e.g. Chakroun et al., 2020; Daw et al., 2006; Wilson & Collins, 2019, Brands et al.,2025).

### Computational Models

Computational model variants differed with regard to their implementation of value updating (i.e. learning) and directed exploration effects.

We compared models based on a classic TD learning algorithm SARSA (QL;Rummery & Niranjan, 1994), such that Q-values of each option *i* on trial *t* are updated according to:

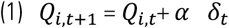

with the reward prediction error (RPE):

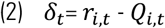

and a constant learning rate *α* (ranging from 0 to 1).

The alternative learning component included uncertainty-dependent updates, which change on a trial-wise basis using a Kalman Filter (Kalman, 1960):

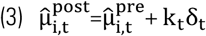

Here 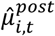 denotes the posterior (i.e. updated) belief about the reward of option *i*, which is updated using the prior belief 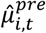 and RPE *δ*_*t*_, scaled by the Kalman gain *k*_*t*_, with:

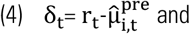

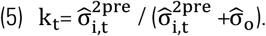

As can be seen in Eq. 5 ,the Kalman gain depends on two different sources of variance, 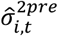 depicts the variance associated with the reward value of option *i*, while 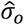 is the observation variance. This parameter indicates the noisiness of observations (vs. the variation in the true underlying reward distribution). For a more exhaustive account, we refer the reader to the preregistration and previous work (see e.g. Chakroun et al., 2020; Daw et al., 2006; Sutton & Barto, 2018).

These two learning mechanisms (QL & BL) were combined with different variants of a basic SoftMax function to yield action probabilities. In the two base models (simply denoted QL & BL) choice probabilities were modelled according to:

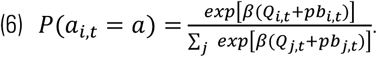

Note that for readability *Q*(*a*) is used for both learning mechanisms, with 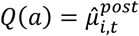 for the BL model variants.

In order to investigate and compare different exploration mechanisms, we tested three variants of an uncertainty bonus. The *classic* implementation assumes that participants update an internal belief model which mirrors the underlying Gaussian random walks (for more details see preregistration; see also e.g. Daw et al., 2006; Chakroun et al., 2020). This version was only combined with the BL base model, as the QL model does not assume participants to represent and update a full environment model (as is the case in Bayesian accounts). This yielded model *BL + Sigma* (c.f. Table 1), where the *exploration bonus* (eb_*i*,*t*_) simply consists of subjects’ (model-based) variance estimates (i.e. beliefs) 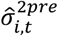 scaled by a free directed exploration parameter ϕ, such that:

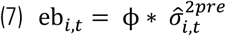

In addition, we also considered two different counter-based uncertainty proxies (see Brands et al., 2025) which are computationally less costly and may therefore be more parsimonious. In one variant (“trial heuristic”) we assume participants to simply track, for each bandit, the number of trials since these have last been sampled, yielding uncertainty proxy 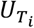 , implemented in models QL + Trial and BL + Trial (c.f. Table 1). In another variant (“bandit heuristic”), we assumed participants to track the number of alternative options that have been sampled since an option was last selected (i.e. ranging from 0 to 3), yielding uncertainty proxy 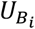 , implemented in models QL + Bandit and BL + Bandit (c.f. Table 1). The resulting exploration bonus for either heuristic was defined analogous to the classic version shown in Eq. 7 above:

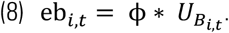

In all models we also accounted for simple perseveration behaviour, defined as repeating the choice from the preceding trial (see also Chakroun et al., 2020). To this end we defined a *perseveration bonus* as:

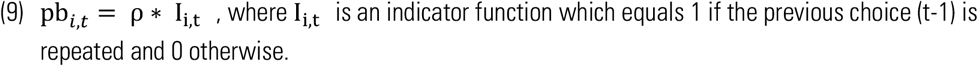

For all models apart from the base versions (c.f. Eq. 6 above) choice probabilities were modelled according to:

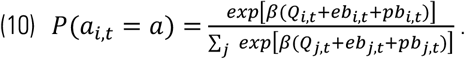

### Exploratory Modelling

In addition to the preregistered model variants, we considered one further adaptation of the previously described SoftMax model, as well as a Drift Diffusion Model (DDM).

#### Higher Order Perseveration

For the SoftMax extension, we incorporated a Higher Order Perseveration (HOP) term, replacing the Fist-Order (FOP) analogue described above. HOP assumes that current choices are not only influenced by the just preceding one (i.e. FOP), but instead are biased by the subject’s choice history reaching further back in time. This process is formalized in the form of a record that stores prior choices independent of any (subjective) reward-values assigned to them (Miller et al., 2019; Palminteri, 2023).

This *habit vector* H_*t*_ is, analogous to subjective reward values of choice options, updated using a decay rate (since we are looking back instead of planning ahead – but can be seen as akin to learning rate parameters). Trial-wise updates of habit strength H_*i*,*t*_ for option *i* on trial *t* follow:

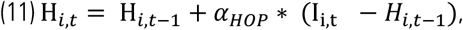

where I_i,t_ is the same indicator function used in the previous FOP model variants (I_i,t_ equals 1 if a choice is repeated and 0 otherwise; c.f. Eq. 9 above). The decay-rate *α*_*HOP*_ updates the habit strength associated with each choice option and determines the extent of the temporal integration window (see also Miller et al., 2019).

Action probabilities follow the same SoftMax as in Equation 10 above, with the only difference that *pb*_*i*,*t*_ is defined as:

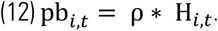

#### Drift Diffusion Model

Next to the models using the SoftMax function to model action probabilities, here we also implemented a Drift Diffusion Model (DDM). As DDMs depict the decision process as a noisy accumulation of evidence towards one of two decision boundaries, we dichotomized choices from the four-armed bandit task by coding choices as either repetitions (*stay*) or alterations (*switch*, Houser, 2025).

Due to their wide applicability and adaptability DDMs have been of great utility to several research foci, including the study of disease-relevant cognitive processing. Aberrant evidence accumulation has thus, been proposed as another candidate for a transdiagnostic neurocomputational phenotype and risk factor (Sripada & Weigard, 2021). Transcending the realm of clinical applications, several research groups have successfully refined, adapted combined this basic DDM architecture with other common computational models e.g. of intertemporal choice (Mathar et al., 2022; Peters & D’Esposito, 2020; Wagner et al., 2020) or as the decision rule in dual-choice RL tasks (e.g. Chakroun et al., 2023; Mathar et al., 2022; Pedersen et al., 2017).

For the current study, and as this is an exploratory addition to our more exhaustive/extensive computational modelling approach presented above, we have limited the DDM-analyses of choice and RT data from the 4-armed restless bandit task on the basic version (i.e. without modifications accounting for learning and subjective valuation).

Basic versions of this model commonly encompass the following key components involved in the decision-process prior to border crossing: the *drift rate* (*δ*), which as the name suggests is a parameter indicating the speed by which evidence is accumulated and thus is negatively associated with the overall RT. The *boundary separation* (*α*) parameter can be interpreted in light of response caution as it regulates the amount of evidence needed until a choice is made (the decision criterion; so that larger values have a prolonging effect of overall RTs; Ratcliff & McKoon, 2008; Wagenmakers, 2009). The parameter *z* determines a potential a priori bias. If the two possible decisions (as was done here) are defined as [0 1], *z*=0.5 would denounce a neutral starting point, while values closer to either boundary denote an according bias (i.e. here values <0.5 would indicate a bias toward choice repetition and >0.5 would indicate one toward switching). Typically, DDMs further account for *non-decision time* with parameter *τ*, which depicts not further specified (e.g. perceptual and motoric) processes that also contribute to response latencies.

In addition to the preregistered RT-preprocessing (exclusion of upper & lower 1% on the group-level and 2.5% on the subject-level), we further set an absolute lower cutoff of 150ms for DDM fitting.

Using the *RWiener* package (Waberisch & Vandekerckhove, 2014) RTs were modelled according to:

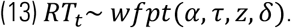

Here the *Wiener First Passage Time* (*wfpt*) describes trial-wise RTs depending on the subject-specific boundary separation *α*, non-decision time *τ*, an initial bias *z*, and the drift rate *δ*.

### Hierarchical Bayesian Modelling in STAN

All models were formulated in the STAN modeling language and were fit using the package *RStan* using the Rstudio interface (RStudio Team 2020, STAN Development Team, 2023). Each model was fit using Markov-Chain-Monte-Carlo (MCMC) via the no-U-turn sampler (NUTS), with four chains running 10000 iterations each, 8000 of which were discarded as warm-up. 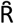 is a measure of convergence across chains, indicating the ratio of between-chain to within-chain variance. Here, values of 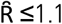 were considered acceptable.

Subject-level parameters were assumed to stem from a shared group-level Gaussian distribution, so that for each model parameter x, the hierarchical models also contained hyperparameters M_x_ and SD_x_ modeling group-level means and standard deviations. For all M_x_ and SD_x_, we set low informative priors assuming them to follow a normal (M=0, SD=10) and uniform (Min=0, Max=20) distribution, respectively. Subject-level parameters were set to be drawn from this resulting group-level normal distribution (M_x_ and SD_x_). All parameters with constraints [0,1] were estimated in standard normal space, and back-transformed to their original range within the model.

Prior distributions for the DDM were likewise low informative and are provided in the supplement (Table S4).

## Results

### Model-Free Results

Descriptive statistics of model-agnostic performance indices separated by factors gender (female vs. male) and condition (PLC vs. TYR) are shown in Figure 2 (see supplemental Table S2). Generally, participants performed above chance, choosing the option associated with the highest reward in 66.57% (SD=6.39; Figure 2A) of the trials, which earned them an average payout of 19000 points per session (SD= 1033; i.e. around 63.36 points per trial). Choice data also contained evidence for exploration behavior, as participants switched to a different option in around 30% of trials (Figure 2B). Median RTs were around .3 seconds (Figure 2C).

**Figure 2.**
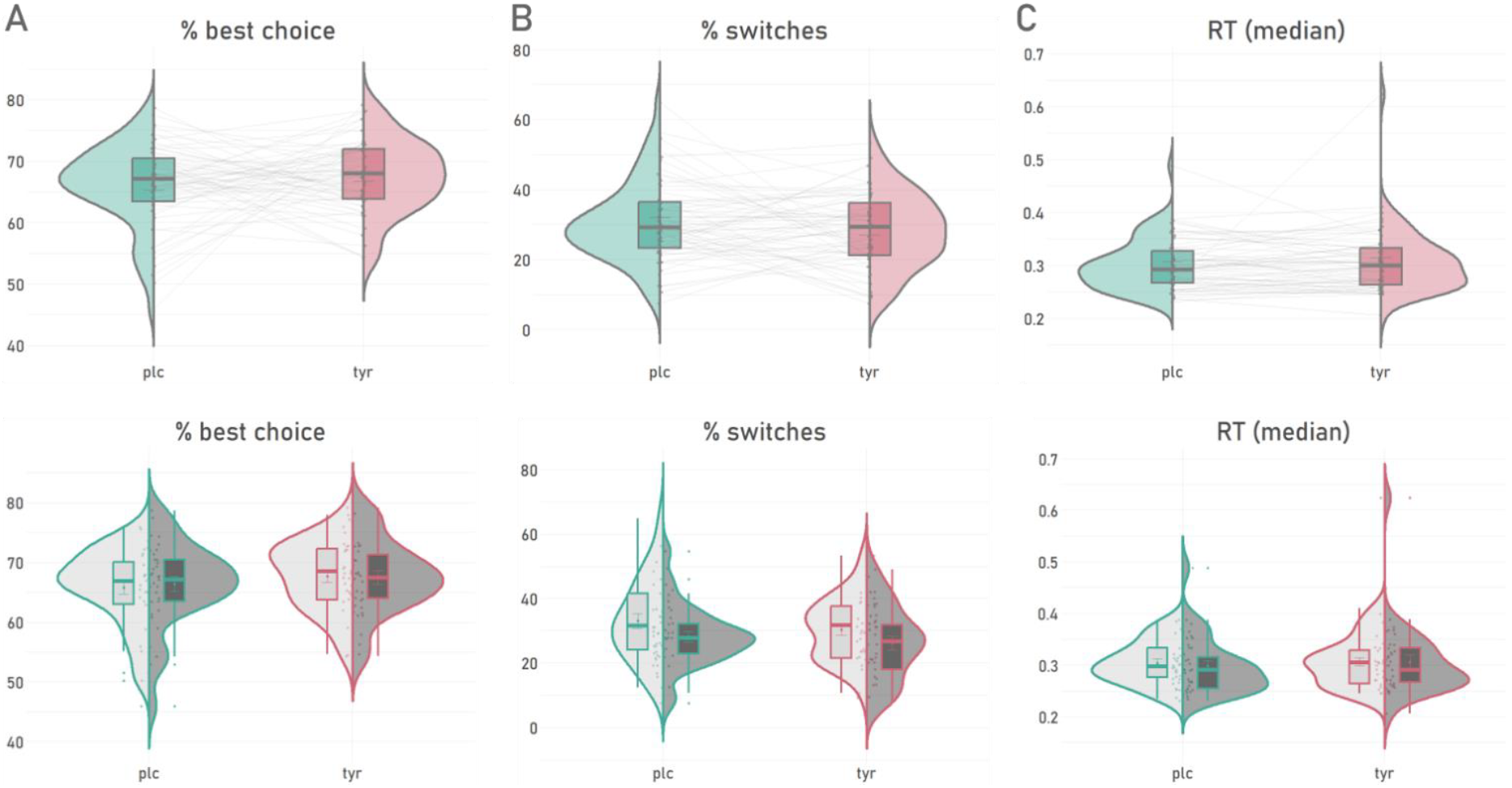
Model-agnostic performance measures separated by supplementation condition and gender group. Supplementation conditions PLC vs TYR are shown in turquoise and magenta, respectively. Dots depict individual subjects. Shading in the lower panel plots mark gender groups (light = female; dark = male). A: percent of choices for the highest-value option. B: proportion of choice alterations (vs. repetitions). C: median reaction times in sec.

The percentage of optimal choices (i.e. choice of the highest rewarded bandit) was increased under TYR vs. PLC) (Table 2; Figure 2A ,main effect of condition p=.019), whereas the condition x gender interaction was not significant (Table 2 ,p>.5). The proportion of switches was significantly reduced under TYR (Table 2 ,Figure 2B ,main effect of condition p<.001), and lower in men vs. women (Table 2 ,main effect of gender p=.029). The regression on median RTs also revealed a small increase in RTs under TYR (see Table 2) that was in the opposite direction compared to previous RT findings in two other tasks (see Mathar et al., 2022).

**Table 2.** Results from Regression Analyses of Model-Free Performance Measures.

|  | *%best choice* |  |  | *RT* |  |  | *%switches* |  |  |
| --- | --- | --- | --- | --- | --- | --- | --- | --- | --- |
| Predictors | Estimate | CI | p | Estimate | CI | p | Estimate | CI | p |
| (Intercept) | 0.66 *** | 0.65 – 0.67 | <0.001 | 0.34 *** | 0.32 – 0.36 | <0.001 | 0.33 *** | 0.30 – 0.36 | <0.001 |
| cond [tyr] | 0.02 * | 0.00 – 0.03 | 0.019 | 0.00 * | 0.00 – 0.01 | 0.045 | -0.02 *** | -0.04 – -0.01 | <0.001 |
| sex [mal] | 0.00 | -0.02 – 0.02 | 0.814 | -0.00 | -0.03 – 0.02 | 0.838 | -0.05 * | -0.09 – -0.00 | 0.029 |
| cond [tyr] *<br>sex [mal] | -0.00 | -0.02 – 0.01 | 0.640 | 0.00 | -0.00 – 0.01 | 0.449 | 0.00 | -0.01 – 0.02 | 0.623 |

\* $p<0.05$ \*\* $p<0.01$ \*\*\* $p<0.001$ ; cond = condition (plc vs. tyr); sex = self-identified gender (female vs. male)

### Model-Based Analyses

Results from the Bayesian hierarchical modelling approach are structured according to the preregistered general modelling procedure.

#### Step 1: Model Comparison & Selection

Using the *loo* package (Vehtari et al., 2017) we first calculated the estimated log pointwise predictive density (elpd), which provides an index for the predictive accuracy of a model via leave-one-out cross-validation. Lower elpd values reflect a better model fit. The superior model in a given set of candidate models therefore has an elpd difference value of 0, and all competitor models are then ranked according to their distance (in elpd) to the best-fitting model (lower elpd_diff_ values indicate a higher relative ranking).

Following our preregistration, model comparison was performed separately for each supplementation condition. This was done to clarify whether the manipulation has an influence on model ranking, which was not the case (c.f. Table 3). The BL+Bandit model provided the best fit to the data across both conditions (see Table 3) and was therefore used for the in-depth analysis of model performance and condition effects on parameters.

**Table 3.**
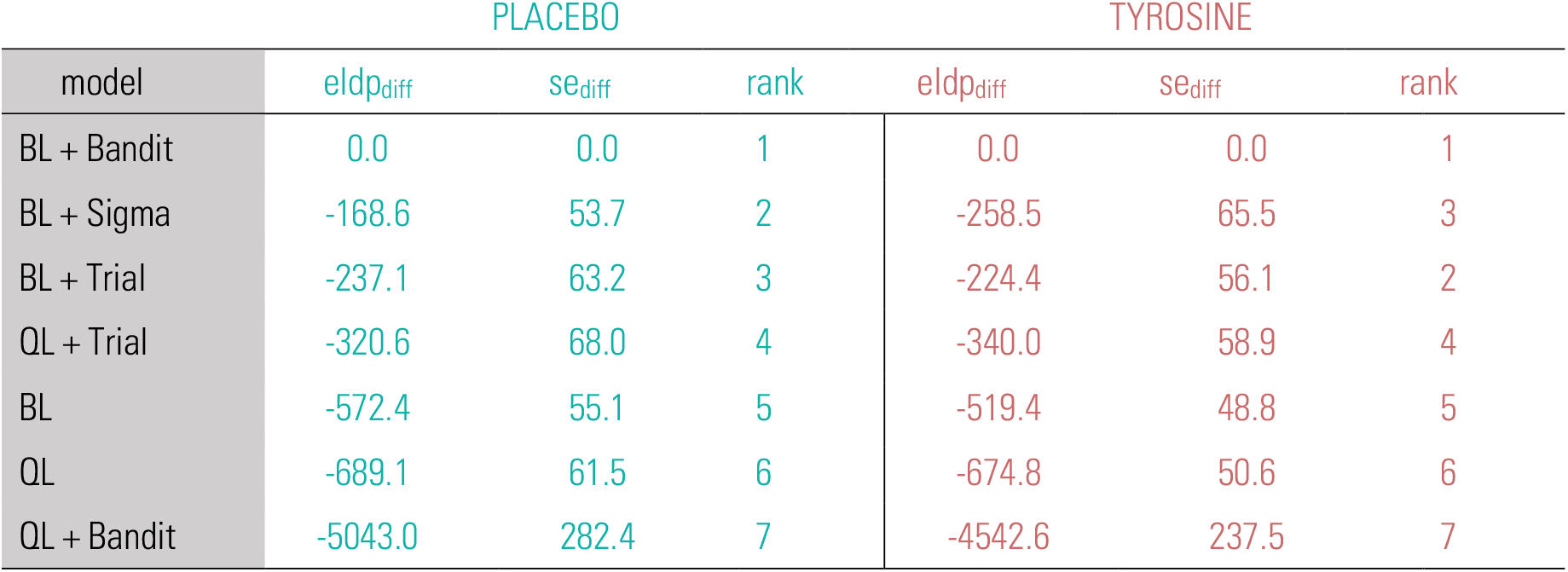
Model Comparison Results Using Leave-One-Out Cross-Validation. BL = Bayesian Learner; QL = Q-Learner; turquoise = PLC; magenta = TYR; elpd_diff_ = difference in estimated log pointwise predictive density; se_diff_ = standard error of the difference.

The posterior distributions of group-level parameters derived from fitting data from the PLC and TYR condition and corresponding summary statistics of the difference distributions (TYR-PLC) are provided in the supplement (Figure S1 and Table S3).

#### Step 2: Modelling of Supplementation Effects

We next examined a combined model that directly included condition-dependent changes in each parameter due to the supplementation. Here, for each parameter, the PLC condition was modeled as the baseline, and changes in each parameter from PLC to TYR were modelled via additional parameters modeling condition effects (“shift” parameters). Thus, resulting posteriors reflect the *shift* in each parameter from PLC to TYR and directly reflect the effect size in units of that parameter.

Parameters reflecting directed exploration (Figure 4B) and perseveration (Figure 4C) exhibited an estimated change largely centered around 0, reflecting the fact that these choice mechanisms were largely unaffected by TYR supplementation. In contrast, the beta parameter reflecting choice stochasticity (i.e. exploitation) was markedly increased under TYR (Figure 4A; 95%HDI≥ 0, see Table 4). This dovetails with the model-agnostic results presented earlier: The increase in choices for the highest value bandit and the decrease in switching behavior can both be attributed to an increase in choice consistency (i.e. an increase in exploitation) under TYR. In contrast, there was no credible evidence for a TYR modulation of directed exploration or perseveration.

**Table 4.** Summary of TYR-Effects on Model Parameters.

| Parameter | Median | 95% HDI | dBF>0 [dBF<0] |
| --- | --- | --- | --- |
| $\beta_{\text{shift}}$ | 0.01 | [ 0.00, 0.02] | 16.94 [0.06] |
| $\Phi_{\text{shift}}$ | -0.09 | [-0.45, 0.34] | 0.52 [1.91] |
| $\rho_{\text{shift}}$ | -0.02 | [-1.32, 1.23] | 0.97 [1.03] |

**Figure 4.**
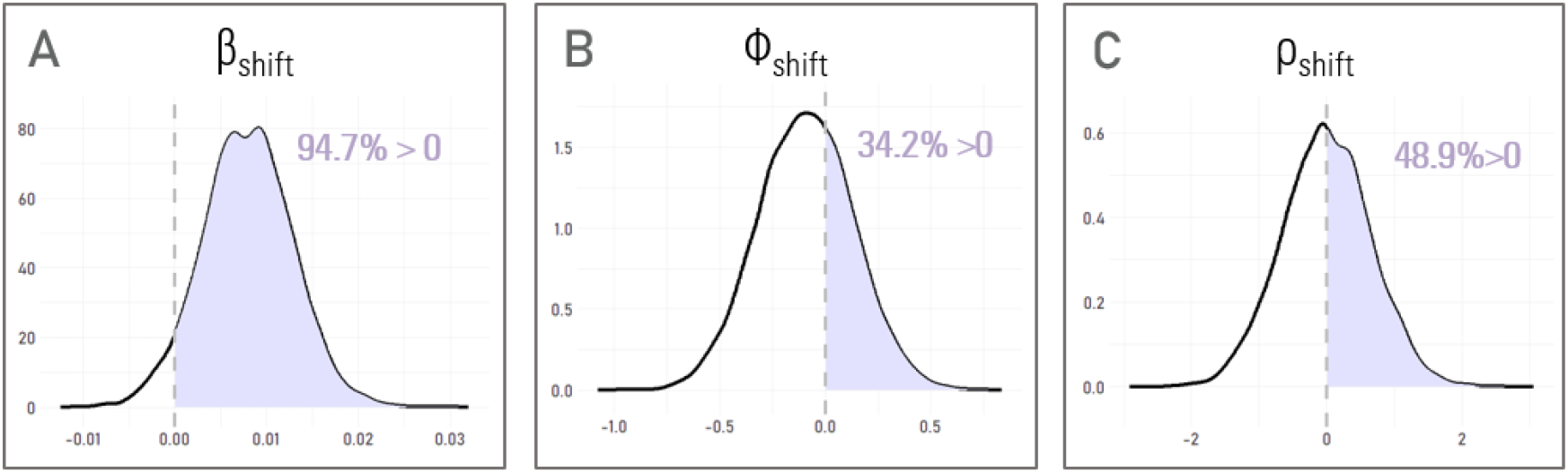
Posterior Distributions of Shift-Parameters Depicting Supplementation Effects. ß: shift (i.e. change) parameter of the SoftMax parameter for value-based choices (i.e. exploitation). Φ: shift of the parameter depicting directed exploration; ρ: shift of the parameter depicting perseveration (choice repetition from t-1). Dashed lines mark a value of 0; percentages show the proportion of the posterior estimates above 0.

To complete the current modelling step, we examined the association of TYR-related changes reported in the section on model-agnostic results with the “exploitation” parameter ß. Correlation analyses (see supplemental Figure S2) confirmed that the TYR-induced increase in exploitation (ß) showed the predicted associations with model-agnostic behavioral measures. Higher exploitation was associated with a lower switch rate, and a greater proportion of optimal choices, in both genders.

#### Step 3: Investigation of gender differences

We next leveraged our comparatively large and gender-balanced sample to examine potential gender-differences in TYR effects. Due to the sparse research on gender-specific behavioral and neurocomputational changes under TYR, we did not formulate specific *a priori* assumptions on these effects. Here, we applied a *shift* version of the best-fitting model (c.f. Step 2) separately to data from self-identified male and female participants. To examine overall gender effects, we first examined the baseline (PLC condition) parameters per gender (Figure 5). There was no credible evidence for gender effects on any parameter (Figure 5 A-C, Table 5). This was further confirmed by a paralleled lack of gender differences in TYR-effects on these parameters (Figure 5 D-F).

**Figure 5.**
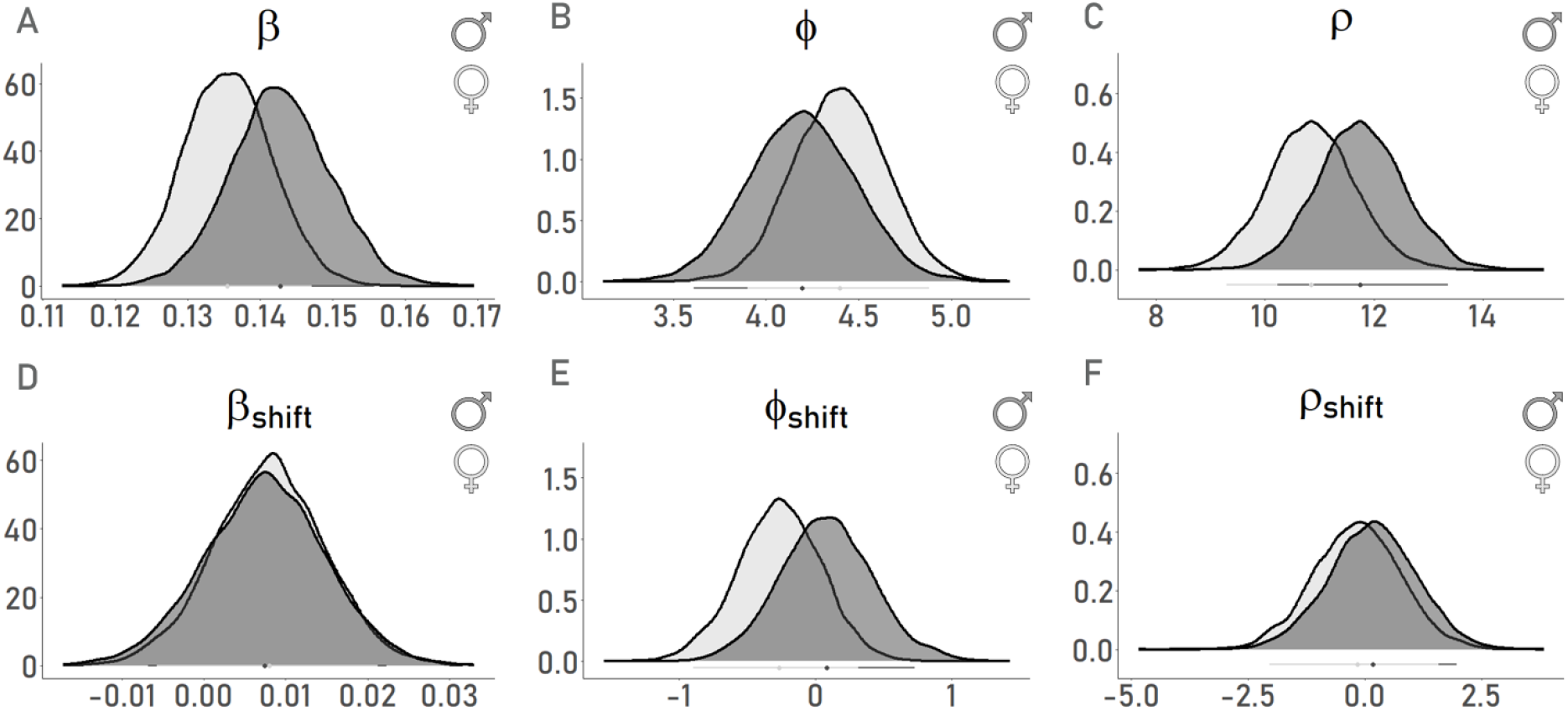
Posterior Distributions for the PLC condition separated by self-identified gender group. A-C: ß: parameter for value-based choices (i.e. exploitation). Φ: directed exploration; ρ: (first-order) perseveration; D-E: TYR-effects on respective model parameters. light and dark gray:results from female and male subsample respectively; bars below the distributions depict the 95% HDI for each group. X-Axis: value estimate of parameter; y-axis: density of the posterior distribution.

**Table 5.**
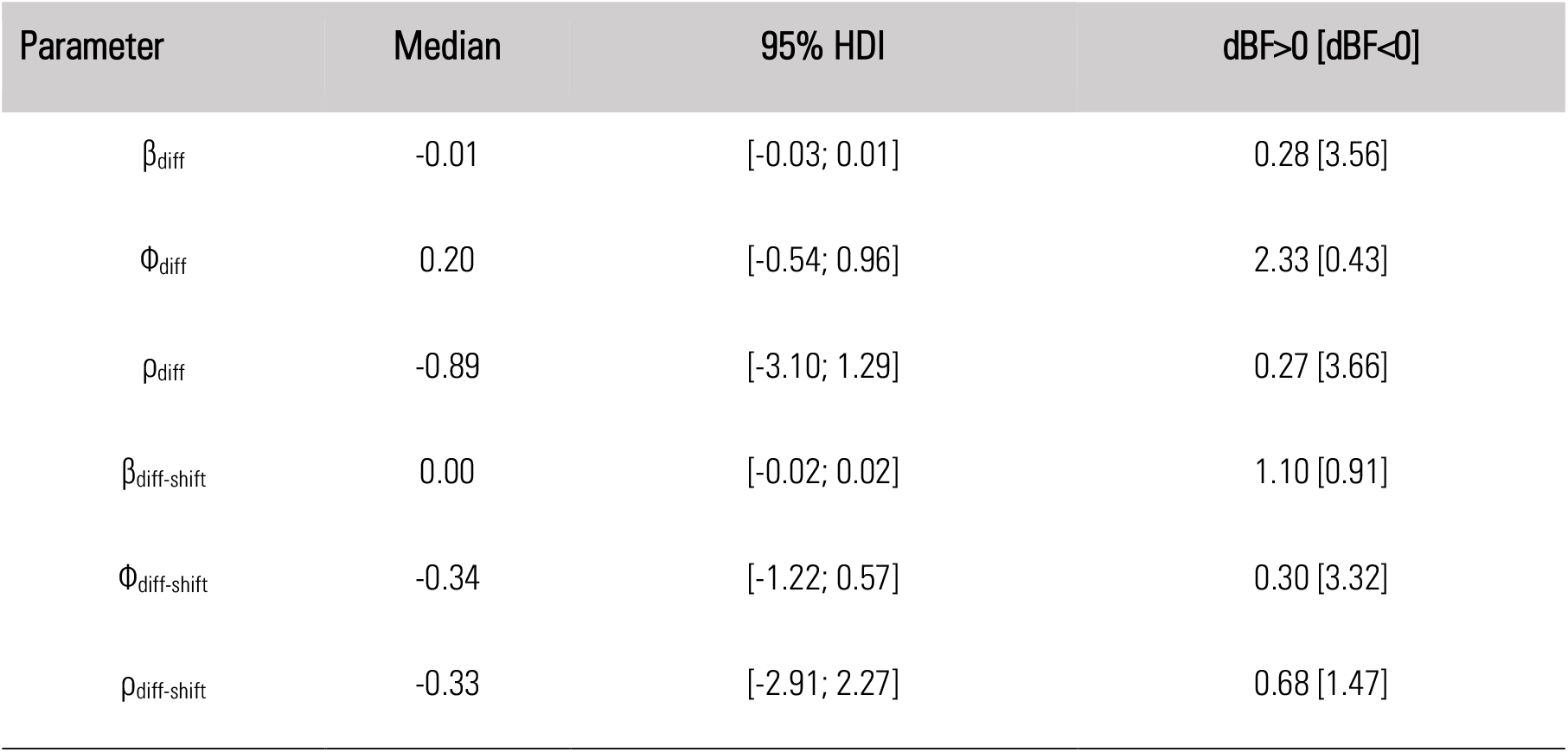
Summary of The Posterior Difference-Distribution Between Gender Groups. Parameters denote the group-level means’ difference distribution (fem-mal) of corresponding SoftMax parameters. dBF>0 [dBF<0]: directional Bayes Factor, calculated as the ratio between posterior density areas over integrals 0:+inf/-inf:0 [-inf:0 /0:+inf].

| Parameter | Median | 95% HDI | $\text{dBF} > 0$ [$\text{dBF} < 0$] |
| --- | --- | --- | --- |
| $\beta_{\text{diff}}$ | -0.01 | [-0.03; 0.01] | 0.28 [3.56] |
| $\Phi_{\text{diff}}$ | 0.20 | [-0.54; 0.96] | 2.33 [0.43] |
| $\rho_{\text{diff}}$ | -0.89 | [-3.10; 1.29] | 0.27 [3.66] |
| $\beta_{\text{diff-shift}}$ | 0.00 | [-0.02; 0.02] | 1.10 [0.91] |
| $\Phi_{\text{diff-shift}}$ | -0.34 | [-1.22; 0.57] | 0.30 [3.32] |
| $\rho_{\text{diff-shift}}$ | -0.33 | [-2.91; 2.27] | 0.68 [1.47] |

**Figure 5.**
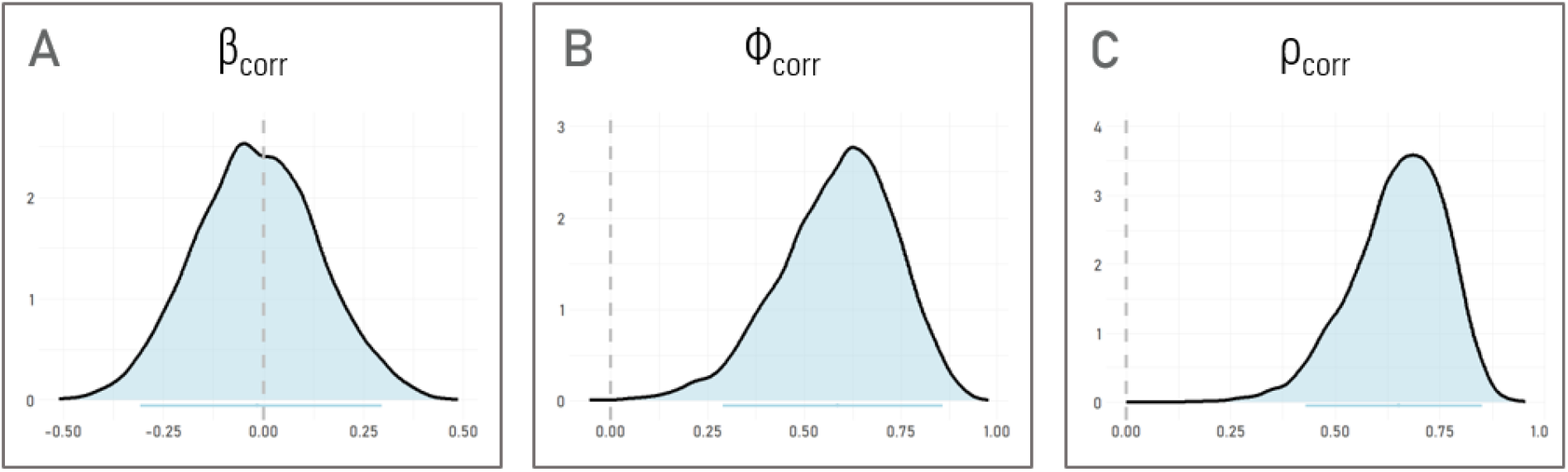
Posterior Distributions of Bayesian Test-Retest Reliabilities. Group-level posterior of the correlation matrices of multivariate model variant (Step 4), showing the test-retest reliabilities (PLC vs. TYR) A: β_corr_, value-based choices (i.e. exploitation); B: Φ_corr_, directed exploration; C: ρ_corr_, perseveration; dashed lines mark x=0; bars below the distributions depict the 95% HDI; x-axis: value estimate of parameter; y-axis: density of the posterior distribution.

Nonetheless, some patterns in the data deserve mention. Numerically, compared to the male sample, female participants showed less exploitation (*stickiness*) and more exploration. These differences may in combination have contributed to the significant gender effect we observed in the model-free analysis of switching behavior (c.f. section Model-Free Results; Table 2; Figure 2).

#### Step 4: Reliability of Parameter Estimates (Multivariate Priors)

Leveraging the comparatively large sample size, we next re-formalized the best-fitting model to obtain a Bayesian estimate of test-retest reliabilities on all model parameters. The hierarchical model was adjusted such that parameters were drawn from bivariate Gaussian Distributions across the two supplementation conditions, allowing us to obtain Bayesian estimated of parameter covariance matrices to compute test-retest reliabilities of model parameters. Figure 5 shows the posterior estimates of test-retest correlations for exploitation (β), directed exploration (Φ), and perseveration (ρ). Group-level means for Φ and ρ exhibited high consistency across conditions (r(Φ)=0.59, r(ρ)=0.65), in line with the finding that these parameters were largely unaffected by supplementation. For exploitation (β), on the other hand, the test-retest reliability posterior distribution was symmetrically centered around 0 (r(β)=-0.02), consistent with a considerable change due to the TYR supplementation. More generally, these findings are in line with the idea that exploitation (β) predominantly reflects state-dependent effects, whereas directed exploration (Φ) and perseveration (ρ) also show considerable within-subject stability.

#### Step 5: Posterior Predictive Checks

In a final step, we examined performance of the best-fitting model using posterior predictive checks by simulating choice data based on draws from each participant’s posterior distribution. We deviated slightly from the preregistered approach (which outlined simulating 500 artificial data sets based on the joint posterior group-level parameters) and examined 8000 simulated data sets per subject using the best-fitting model BL + Bandit.

Each simulated data set was based on a subject-specific parameter combination drawn from the participants’ posterior distributions. Each simulated data set was compared to the respective observed data. Median correct choice predictions separated by supplementation condition are shown in Figure 6A. Prediction accuracy was at a similar level for both conditions and well above chance level (PLC: Mdn=66.31%; TYR: Mdn=67.54%). We next evaluated, for each participant, each of the 8000 full simulated data sets using model-agnostic measures (% best choice, %switches) and averaged these predicted scores across simulations. These largely reproduced the observed behavioral data in both conditions and both for % best choices (Figure 6B) and switch rate (Figure 6C), although the latter was slightly overestimated by the model. Taken together, posterior predictive checks confirmed that the best-fitting model reproduced key patterns in the data.

**Figure 6.**
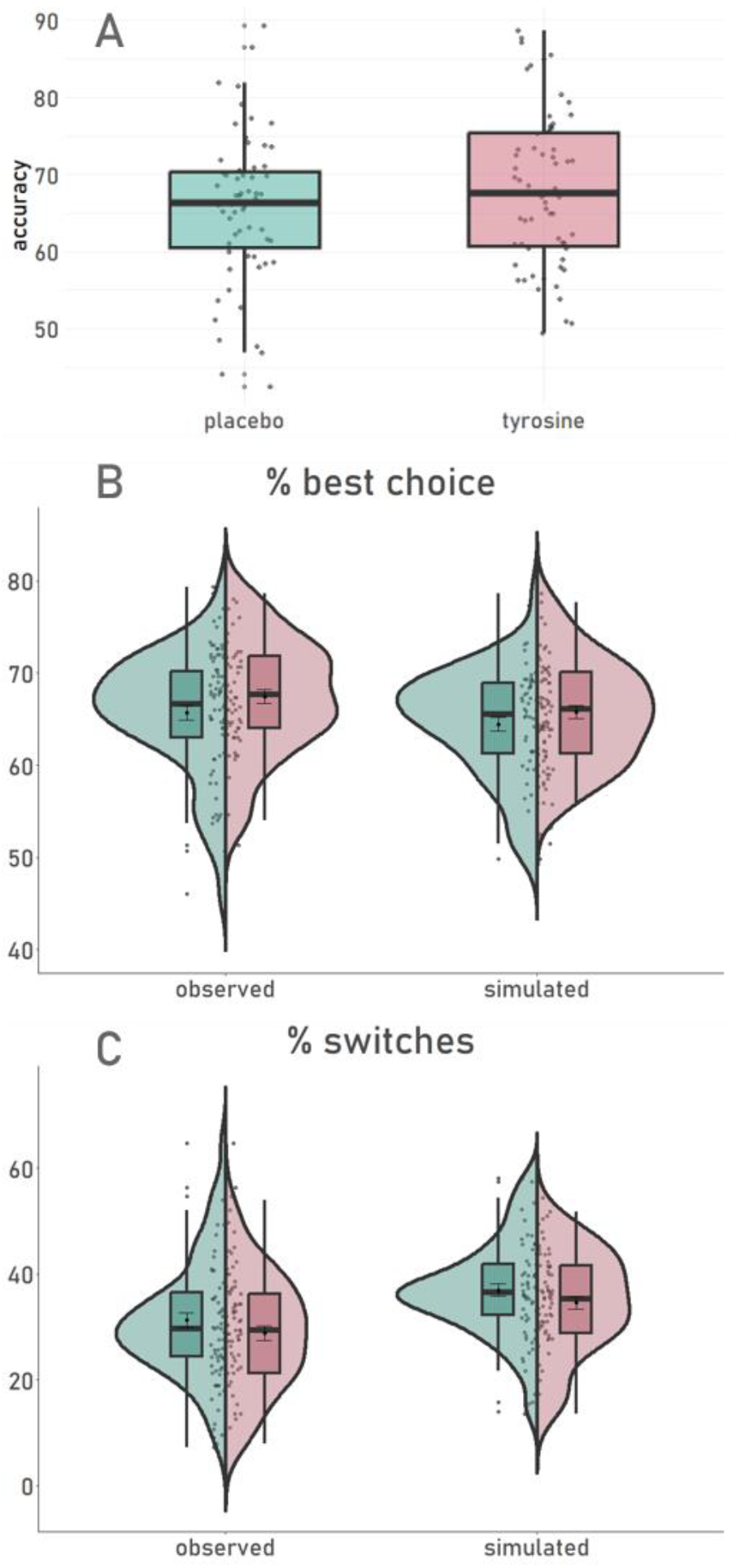
Analysis of Simulated vs. Observed Choice Data. Turquoise = PLC, Magenta =TYR, Points represent participants. A: predictive accuracy calculated as the proportion of correct choice predictions by the winning model compared to observed choices. Panel B & C: Proportion of highest-value choices (%best) and choice alterations (%switches) separated by condition. Observed = empirical data (c.f. model-agnostic results); simulated = predicted choice data based on the winning model.

#### Step 6: Exploratory Modelling

##### 6.A. Higher Order Perseveration

We ran additional, non-preregistered modelling analyses, based on previous results that perseveration behavior may extend across longer choice sequences (higher-order perseveration; Miller et al., 2019, Brands et al. 2025; Tuzsus et al. 2024, Palminteri, 2023). As mentioned above, we consequently set up another refined version of model BL + Bandit in which the indicator function depicting FOP was replaced by the mechanisms suggested by Miller et al (2019), i.e. a *habit vector* H_*t*_ which tracks the choice history for each bandit (Gershman, 2020; Miller et al., 2019) (see Methods section).

This extended model (*Habit;* Table 6) was only applied to the previous BL model variants, which outperformed all QL models; (c.f. Table 2), and we focused on the base model and exploration models BL+Bandit & BL+Sigma. Model comparison favored model variants accounting for HOP, with the extended BL+Bandit model showing the best fit across both conditions.

**Table 6.** Model Comparison Results via Leave-One-Out Cross-Validation. BL = Bayesian Learner; Habit = HOP extension (vs. FOP); turquoise = PLC; magenta = TYR; elpd_diff_ = difference in estimated log pointwise predictive density; se_diff_ = standard error of the difference.

| model | PLACEBO |  |  | TYROSINE |  |  |
| --- | --- | --- | --- | --- | --- | --- |
| | $\text{elpd}_{\text{diff}}$ | $\text{se}_{\text{diff}}$ | rank | $\text{elpd}_{\text{diff}}$ | $\text{se}_{\text{diff}}$ | rank |
| BL + Bandit + Habit | 0.0 | 0.0 | 1 | 0.0 | 0.0 | 1 |
| BL + Sigma + Habit | -91.3 | 61.6 | 2 | -101.1 | 60.9 | 2 |
| BL + Bandit | -168.9 | 36.3 | 3 | -102.2 | 20.4 | 4 |
| BL + Sigma | -661.8 | 95.5 | 4 | -744.1 | 67.3 | 5 |
| BL + Habit | -691.0 | 59.2 | 5 | -613.9 | 58.1 | 3 |
| BL | -1193.6 | 132.7 | 6 | -980.6 | 77.3 | 6 |

Figure 8 depicts posterior estimates of supplementation effects for the four main parameters of interest (c.f. Results on the BL+Bandit model with the addition of HOP step-size α_HOP_). Modeling results of supplementation effects accounting for HOP reproduced the previously observed increase in exploitation (β) under TYR (Figure 8A and Table 7). However, the extended model also revealed credibly reduced parameters for directed exploration as well as perseveration (Φ and ρ, respectively, 95% HDI < 0; Table 7 ,Figure 8C & D) under TYR. Accounting for HOP therefore led to a more fine-grained decomposition of choice-mechanisms, and now revealed an attenuation in directed exploration and perseveration under TYR as well as substantiating the previously moderate effect on exploitation.

**Table 7.** Posterior Estimates From Step 2 (Shift Model) of the Adapted BL + Bandit + Habit Model. β_shift_= effect on value-based decision making; Φ_shift_= effect on directed exploration; ρ_shift_ = effect on perseveration (HOP); . dBF>0 [dBF<0]: directional Bayes Factor, calculated as the ratio between posterior density AUCs over integrals 0:+inf/-inf:0 [-inf:0 /0:+inf].

| Parameter | Median | 95% HDI | $\text{dBF}>0$ [$\text{dBF}<0$] |
| --- | --- | --- | --- |
| $\beta_{\text{shift}}$ | 0.01 | [ 0.00, 0.02] | 60.75 [0.02] |
| $\Phi_{\text{shift}}$ | -0.48 | [-1.03, 0.00] | 0.03 [31.15] |
| $\rho_{\text{shift}}$ | -1.85 | [-3.64, -0.02] | 0.02 [47.99] |
| $\alpha_{\text{HOP}}\text{-shift}$ | 0.04 | [ 0.00, 0.08] | 45.43 [0.02] |

**Figure 8.**
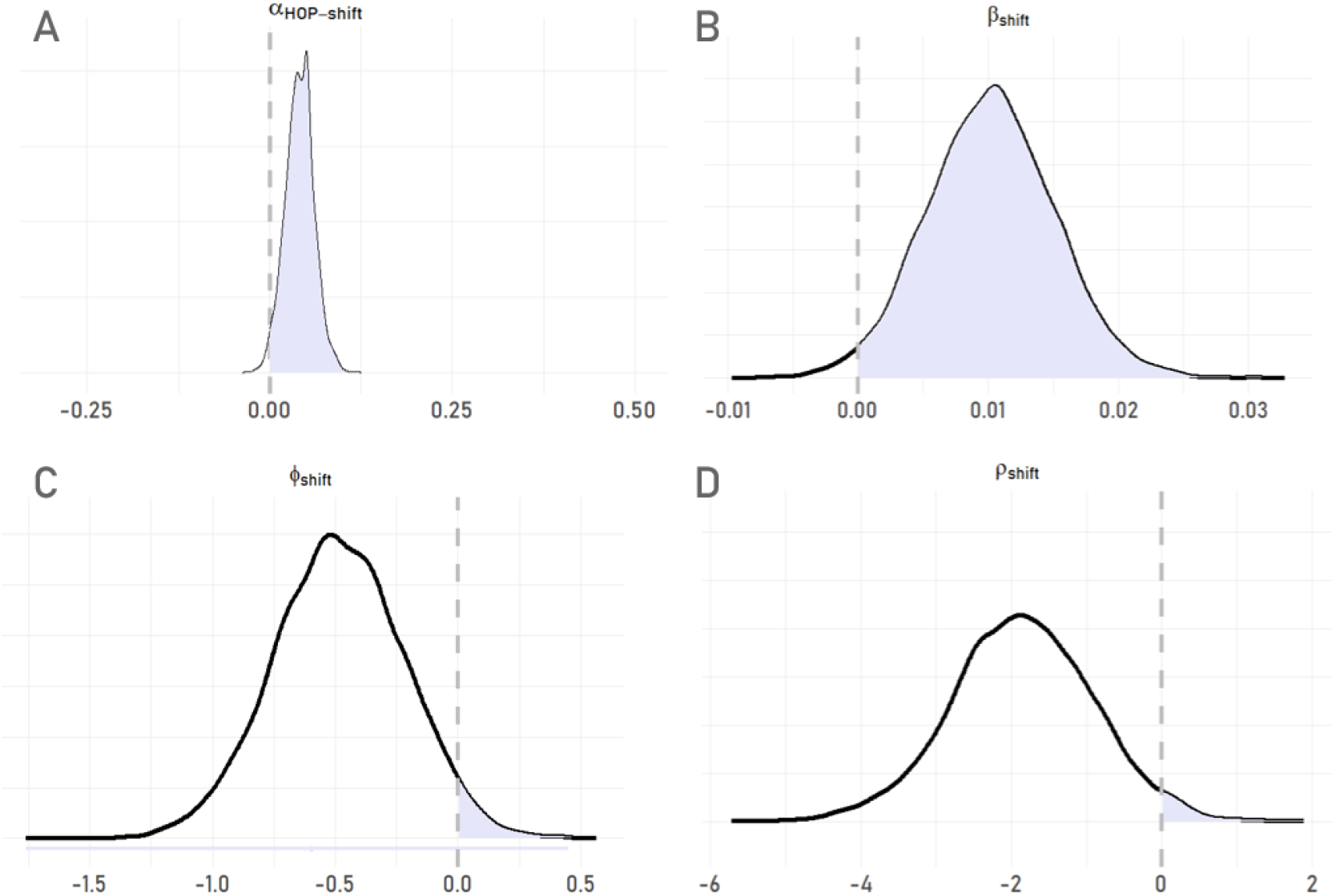
Posterior Distributions of TYR Effects on Parameters From HOP Model-Extension. Depicted are the posterior distributions from Modelling Procedure Step 2 based on the BL+Bandit+Habit Model. “Shift”-Parameters indicate the supplementation effect (PLC vs. TYR) on corresponding parameters: A: higher-order perseveration step-size *α*_*HOP*−*shift*_; B: exploitation *β*_*shift*_; C: exploration *Φ*_*shift*_; B: perseveration *ρ*_*shift*_ . Dashed lines mark a value of 0; shaded areas mark values <0.

Results from fitting the extended model to each condition (Table S5) and gender group separately (Figure S3 ,Table S6) are provided in the supplement.

##### 6.B. DDM: stay vs. switch choices

As preregistered we also conducted a model-based analysis of RT-distributions on the bandit task using a Drift Diffusion Model (DDM). In this model each decision is modeled as a noisy evidence accumulation process toward one of two boundaries. These mark the criterion for a decision, so that crossing either *decision boundary* is equivalent to the observable point at which a choice is made (Meyers et al., 2022; Ratcliff & McKoon, 2008). In order to dichotomize the four possible responses on the task, we coded each choice on trial t as either a stay (0) or switch (1) relative to the previous choice on trial t-1. This FOP-categorization yielded the decision boundaries for the DDM-analysis (c.f. Houser, 2025).

Graphic and tabular depictions of posterior parameter estimates per condition and the resulting difference distributions can be found in the supplement (Figure S4 ,Table S7). Modelling of TYR effects on DDM parameters in a combined model indicates that non-decision time (*τ*_*shift*_) and boundary separation (*α*_*shift*_) were largely unaffected. HDIs of the a priori bias (*z*_*shift*_; 95% HDI=[-0.01, 0.05], dBF>0=13.96) and the rate of evidence accumulation (δ_shift_; 95% HDI=[-0.45, 0.04], dBF<0=17.48), show more noteworthy deviation from 0 while, however, still not providing credible evidence for an effect (Figure 9 ,Table 8).

**Table 8.** Posterior Parameter Estimates Derived from a DDM With Stay-Switch Decision-Boundaries. α = boundary separation; z = bias; δ = drift rate; τ = non-decision time; dBF = directional Bayes Factor.

| Parameter | Median | 95% HDI | $\text{dBF}>0$ [$\text{dBF}<0$] |
| --- | --- | --- | --- |
| $\alpha_{\text{shift}}$ | 0.01 | [-0.02, 0.04] | 3.89 [0.26] |
| $z_{\text{shift}}$ | 0.02 | [-0.01, 0.05] | 13.96 [0.07] |
| $\tau_{\text{shift}}$ | 0.002 | [-0.01, 0.01] | 2.28 [0.44] |
| $\delta_{\text{shift}}$ | -0.20 | [-0.45, 0.04] | 0.06 [17.48] |

**Figure 9.**
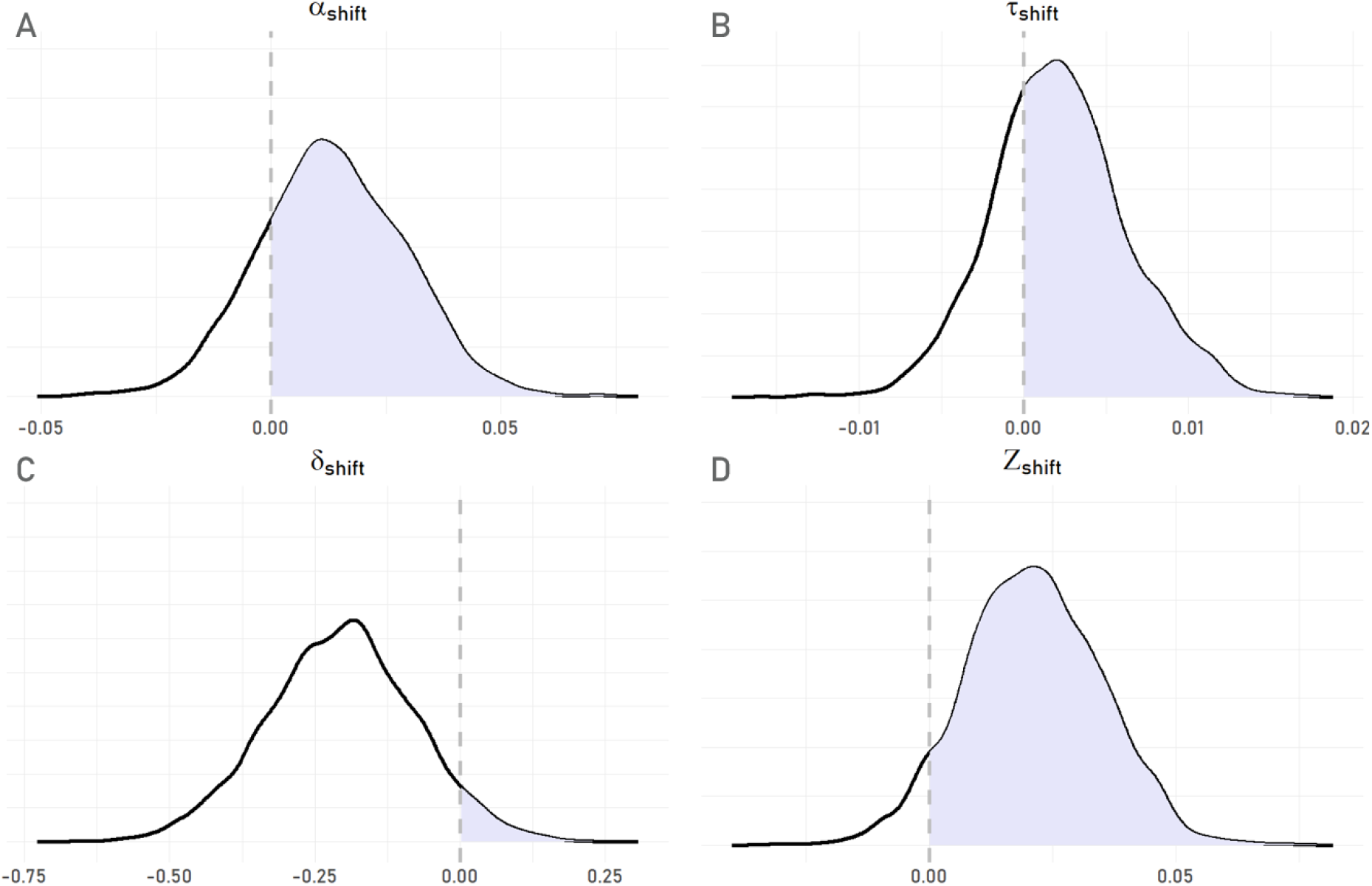
Posterior Distributions of DDM-Parameters Depicting Changes Due To Condition. Group-level shifts in parameter estimates from condition PLC to TYR;A: α = boundary separation; B: τ = non-decision time; C: δ = drift rate; D: z = bias. Decision Boundaries were defined as 0 = stay (i.e. repetition of preceding choice); 1= switch (choice alteration).

The above reported model-free slowing effect of TYR on median RTs thus, may be driven by contrasting effects: a small shift of the starting point in favor of a choice switch (increased *z*, Figure 9D) paired with a simultaneous increase in the drift toward a choice repetition (decreased *δ*, Figure 9C). Recall, however, that this analysis was based on the dichotomization into *stay* and *switch* choices, while the referred to main effect of RT-slowing under TYR did not make this distinction.

Thus, overall examination of the combined model including condition effects on DDM parameters (Figure 9 ,Table 8) revealed no credible evidence for TYR-related effects.

## Discussion

Here we show that a single dose of the catecholamine precursor TYR modulates reinforcement learning and decision-making in a commonly applied RL task with a dynamic reward schedule. Most notably, we found an overall increase in choice consistency and optimal choices under TYR, and both effects were largely independent of gender. Comprehensive computational modelling further revealed that TYR supplementation improved performance by increasing value-based exploitation, in line with the observed increase in optimal choices and a reduction in the switch rate. Extending this model variant by also accounting for higher-order perseveration (HOP) provided a further improvement of model fit, and revealed more fine-grained insights into latent processes. While the increase in value-based exploitation was substantiated in this model, TYR furthermore reduced directed exploration and value-independent perseveration, effects that were potentially masked in the previous, simpler model variant. This was accompanied by a change in HOP step-size, indicating a reduced integration window of prior choices under TYR. These results provide insights into the neurocomputational mechanisms underlying TYR supplementation effects on RL performance.

Model comparison revealed that behavior was best accounted for by a model that assumes an uncertainty-dependent learning process (i.e. Bayesian Learner via Kalman Filter). The best-fitting model further included a heuristic-based directed exploration mechanism (i.e. “Bandit Heuristic”, where participants track how many other options were sampled since a specific option was last selected), as in related work (Brands et cl 2025). These modelling results are in line with previous studies that also favored Kalman Filter over less flexible and uncertainty-naïve Q-learning variants (see e.g. Chakroun et al., 2020; Daw et al., 2006; Wiehler et al., 2021). However, in the present study, directed exploration was better accounted for by a heuristic process, as opposed to the uncertainty estimate of the Kalman Filter model (c.f. e.g. Chakroun et al., 2020; Daw et al., 2006, Fox et al., 2020).

Analysis of this preliminary best-fitting model revealed that the performance improvement under TYR was due to an increase in value-driven exploitation (beta parameter of the softmax function) (Rutledge et al., 2015; Kroemer et al., 2019; Adams et al., 2020). Parameters reflecting (heuristic-based) directed exploration and perseveration (FOP) did not exhibit meaningful variation across conditions based on this model variant, although numerical trends pointed toward a reduction in these two components under TYR. Extending this model by accounting for HOP in an additional exploratory modelling step, however, revealed a substantial improvement in model fit. It also substantiated the previously observed increase in value-based exploitation (softmax beta) under TYR, whereas TYR-related reductions in directed exploration and value-independent perseveration became considerably more substantial. This change due to model refinement echoes previous findings of improved granularity upon defining and formalizing related processes (Mkrtchian et al., 2023; Wilson & Collins, 2019). For example, using the same task employed here, Chakroun et al. (2020) observed an increase in directed exploration estimates after accounting for FOP, compared to a model variant without a perseveration term (Daw et al., 2006). Specifying previously unaccounted-for processes can thus aid the identifiability and interpretability of other model components by accounting for regularity in the data that would otherwise reflect unexplained variation.

The work by Chakroun et al. also relates to the present results with respect to the role of DA, as it involved a more direct manipulation of DA levels via a pharmacological intervention with either Haloperidol or Levodopa (LDOPA; vs. PLC). Based on their computational model, L-DOPA reduced directed exploration compared to both placebo and the D2 antagonist Haloperidol. Although a direct comparison is complicated by the different approaches to catecholamine modulation (pharmacological vs. supplementation-based), these results converge with our observation of reduced directed exploration under TYR in the best-fitting model variant. TYR supplementation presumably elicits a more indirect manipulation of DA-levels compared to pharmacological approaches (Jongkees et al.,2014, 2015; Magill et al., 2003; Bloemendaal et al., 2018; Hase et al., 2015). Nonetheless, the observed reduction in directed exploration observed here dovetails with the results by Chakroun et al. (2020).

Mathar and colleagues (2022) followed the same supplementation regime as done here but examined a different RL-task (two-step task; Daw et al., 2011). Although this complicates a direct comparison of results, it may explain contrasting RT effects of TYR in Mathar et al. and the present study. Mathar et al. observed RT reductions under TYR for the two-step task, whereas we observed RT increases for the bandit task, and this might be related to the specific decision problem. The two-step task is more complex due to its sequential, probabilistic architecture, and involves binary decisions on all stages, whereas participants decide between four options on the bandit task. Demands therefore clearly differ between tasks. Across both studies and tasks, however, participants exhibited a higher tendency to repeat vs. alternate preceding actions following the ingestion of TYR, consistent with the idea of choice stabilization. Generally, TYR effects may manifest differently depending on the task demands and performance-related processes.

In order to provide more complete perspective on the observed data, we also modeled RT-distributions for choice switches vs. repetitions using a variant of the drift diffusion model (DDM).This analysis, independent of further assumptions regarding learning and valuation processes, revealed a supplementation effect on the drift rate parameter, reflecting a tendency for an increased rate of evidence accumulation (in favor of choice-repetitions under TYR. However, an opposing effect was found for the bias (starting point), which was shifted towards the “switch”-boundary. These contrasting effects may partially explain the small yet significant increase in median RTs under TYR (vs. PLC) observed in our model-agnostic analyses.

The present study focused on effects of a single TYR dose, and did not investigate long-term effects of supplementation. While there are first reports indicating positive long-term effects of TYR on domains such as working memory, and fluid intelligence (Kühn et al., 2019), more exhaustive investigations are clearly required. Such future work would preferably also include more diverse samples. Another aspect that would be of interest in the context of longitudinal studies in light of our results (exploratory modeling using HOP) concerns perseveration and habitual responding. Because a key differentiating factor between perseveration and habits concerns their temporal extension, long-term investigations might shed light on TYR effects on perseveration and/or habitual responding over time. While perseveration can to some extent be measured in a cross-sectional laboratory-based studies, habits are much harder to capture as they have developed over and may persist across more extended time periods (Bornstein & Banavar,2023; see also Buabang et al., 2025; Nebe et al, 2024).

There are several potential modes of action of a low dosage supplementation regime as implemented in the present study (see e.g. Jongkees et al., 2015, 2020; Bloemendaal et al., 2018; Hase et al., 2015). A common interpretation of TYR effects is based on the assumption that increasing precursor availability (e.g. via supplementation) can increase neurotransmitter synthesis under demanding conditions (Attiepoe et al., 2015; Jongkees et al., 2014, 2015; Kvetnansky et al, 2009), However, individual supplementation effects may depend on individual differences in the DA system (Pinto & Uchida, 2023), which may contribute to mixed effects in the literature. Furthermore, the degree to which catecholamine neurotransmitters beyond DA may be affected by TYR remains unclear. Even when focusing on the role of DA in RL, as probed by e.g. the restless bandit task applied here, supplementation may affect phasic and/or tonic DA levels (Beeler et al., 2010; Glimcher, 2011; Pinto & Uchida, 2023) and effects may additionally depend on individual factors such as baseline availability and synthesis capacity (see e.g. Jongkees et al., 2014; Hase et al., 2015).

Overall, current findings are in line with recent theories on the role of DA in reinforcement learning and decision-making (Niv et al., 2007; Friston et al., 2012; Mikahel et al., 2021). The theory of *Rational Inattention* for example, posits that precision in learning and reliance of actions on learned representations (e.g. reward magnitudes) heavily rely on individuals’ dopaminergic functioning. Mikahel and colleagues (2021) state that high (vs. low) tonic DA levels foster learning from rewards (vs. punishment or low reward) and influence action control in favour of exploitation (vs. exploration). Constituent mechanisms here may include a more pronounced contrast in subjective values (due to more precise estimates) and a resulting preference for higher value options (i.e. exploitation over exploration) (see also Wagner et al. 2020). Thus, under the assumption that TYR supplementation in the present study modulated DA-tone, our results largely align with these ideas.

### Limitations & Future Directions

Despite the relatively large and gender-balanced sample and comprehensive modeling procedures, several limitations of our approach need to be addressed. Conclusions are limited to western, educated, white and cis-gendered, mostly young and able-bodied volunteers, limiting generalizability. We also did not examine potential dosage effects, but used a widely-used fixed dosage of 2g (see e.g. Jongkees et al., 2015; Steenbergen et al., 2015; Dennison et al., 2019; Robson et al., 2020; Mathar et al., 2022). Recommendations and applications with regard to TYR-doses vary across previous investigations (e.g. Deijen 2005: 14mg/kg), which precludes direct comparison of findings and may contribute to mixed effects. However, control analyses of model-agnostic effects that included weight as an additional random effect confirmed our results (Supplementary Table S8). Nonetheless, even a more direct and rigorous tracking of catecholamine levels would not eliminate variability of TYR ingestion via other exogenous sources, such as dietary choices. While for the current study, we did ask participants to abstain from large, protein-rich meals in advance of testing sessions, adherence to this request could not be directly controlled. It should be noted though, that despite their potential influence on present findings, TYR-related effects found were stable across gender groups as well as age and weight ranges (as narrow as these may have been).

Although our results do provide new insights into latent effects of TYR on cognitive subprocesses, our study design only allowed for a cross-sectional snapshot of such effects. Investigations into long-term effects of TYR (self-) administration on (among others) learning and decision-making are called for (see above). This seems especially important as the practical application of such supplements is not limited to a one-time low-dose ingestion. Indeed, between 40-60% of customers in India (and around 45% in the United States) reported daily use of dietary supplements in a recent survey (Rakuthen Insight, 2022; Statista, 2020).

As outlined previously, the exact neurochemical effects of TYR are still only partially understood, but are thought to be modulated by baseline DA levels (see e.g. prominent related work by Jongkees et al., 2015 on a potential inverted-U shaped association of DA and executive functions). Consequently, effects of additional exogenous manipulation of catecholamine synthesis (e.g. via supplementation) may elicit different effects depending on such endogenous factors.

Another limitation may lie in the analysis approach applied. Despite their clear advantages and benefits compared to model-free compound measures, cognitive computational models have their own inherent limitations. Model variants considered in this project cover a considerable range of established accounts, but all conclusions remain restricted to the examined model space. For example, even the best-fitting model only accounted for around 67% of choices, on average, such that there is clearly room for further improvements. We wish to address and tackle this issue head-on by making our analysis procedure, including computational details along with explicit model codes available. Other interested researchers are invited to test and tweak modelling accounts presented here, and report on their results obtained in this endeavor.

## Conclusion

We investigated the effect of a single dose of the catecholamine precursor tyrosine (2g) on RL and decision-making in a young and healthy gender-balanced sample. Tyrosine improved performance in terms of the percentage of optimal choices, and extensive computational analyses linked this effect to an increase in value-based exploitation. Further model extensions that accounted for higher-order choice perseveration effects (Miller et al., 2019; Brands et al. 2025; Tuzsus et al. 2024, Palminteri, 2023) substantiated this effect, and further clarified underlying mechanisms by revealing a reduction in directed exploration (resonating with previous pharmacological studies, Chakroun et al., 2020) and value-independent perseveration under TYR. In a comparatively large and gender-balanced sample, our results reveal robust effects of TYR on neurocomputational processes thought to be under dopaminergic control.

## SUPPLEMENT

**Table S1.**
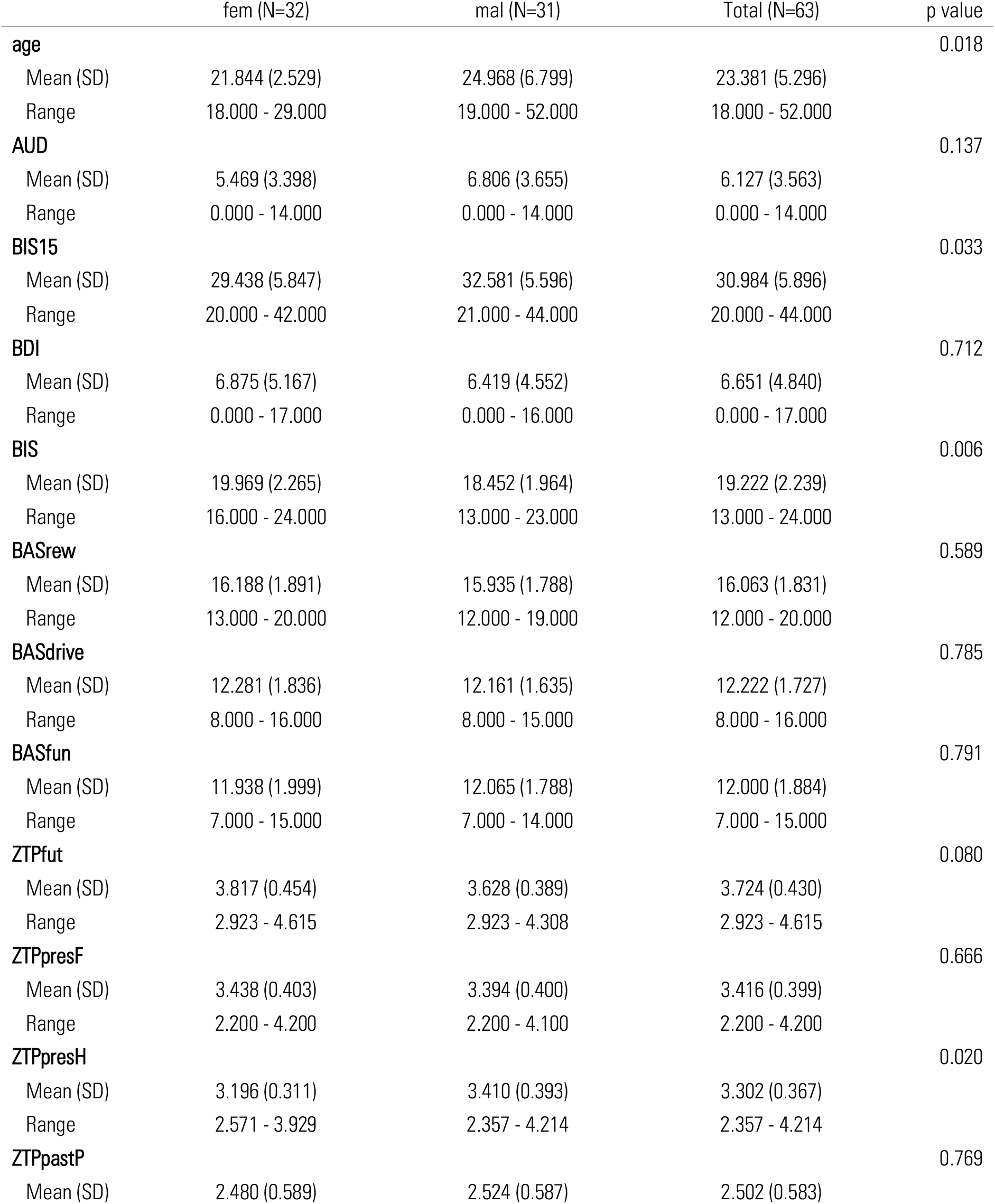
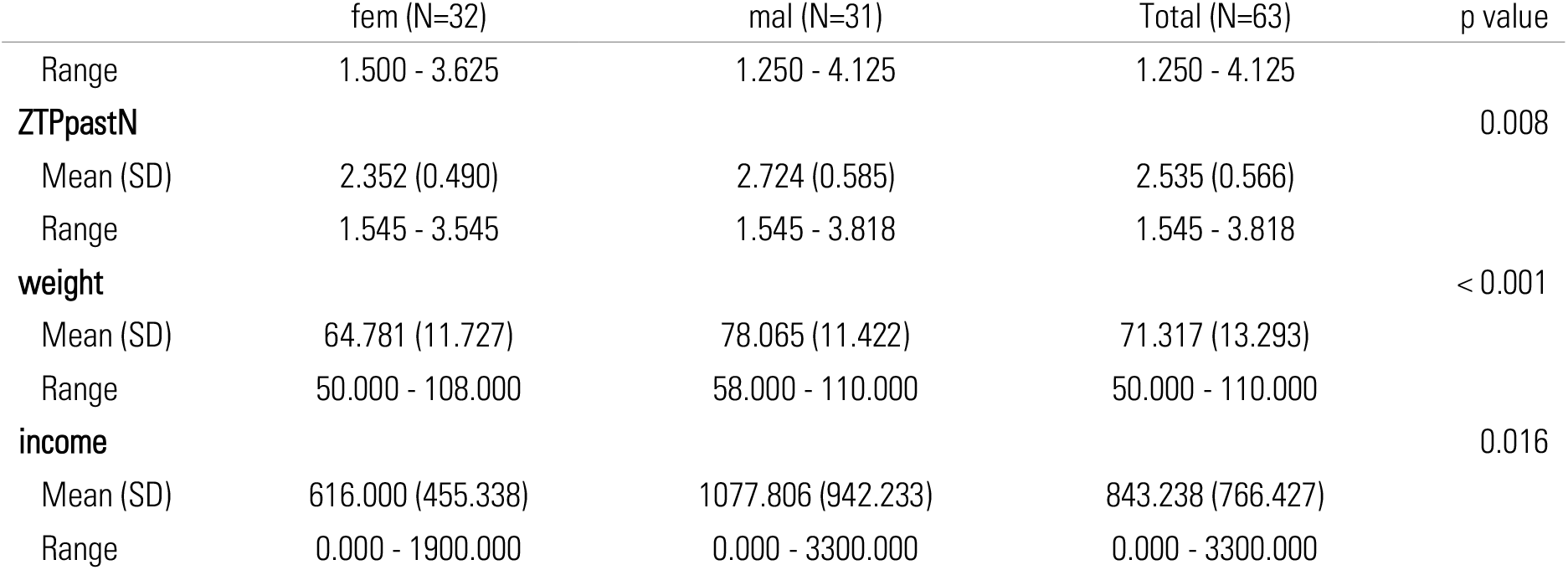
Summary statistics of self-report measures and basic demographic information.

**Table S2.** Summary statistics of model-agnostic performance measures separated by gender group and supplementation condition. Reward: accumulated reward per experimental session. RT(med): median RT. %Best: proportion of choices for the highest reward bandit. %Switch: proportion of choice alterations (vs. repetition). Dem/mal: self-identified male and female gender.

|  | fem:plc (N=32) | fem:tyr (N=32) | mal:plc (N=31) | mal:tyr (N=31) |
| --- | --- | --- | --- | --- |
| **Reward** |  |  |  |  |
| M (SD) | 18956.41 (1050.24) | 18969.11 (1007.10) | 19053.660 (1046.73) | 19053.69 (1074.27) |
| Range | 17165.97- 20420.41 | 17332.84- 20571.23 | 17410.50- 20587.33 | 17612.55- 20561.0 |
| **RT (med)** |  |  |  |  |
| M (SD) | 0.298 (0.047) | 0.303 (0.049) | 0.286 (0.069) | 0.295 (0.084) |
| Range | 0.181 - 0.386 | 0.229 - 0.410 | 0.077 - 0.487 | 0.062 - 0.624 |
| **%Best** |  |  |  |  |
| M (SD) | 65.500 (6.336) | 67.271 (5.992) | 65.882 (6.890) | 67.645 (6.393) |
| Range | 50.667 - 75.667 | 54.000 - 78.000 | 46.000 - 79.333 | 54.333 - 78.667 |
| **%Switch** |  |  |  |  |
| M (SD) | 33.396 (12.393) | 30.854 (9.762) | 29.108 (9.385) | 26.742 (11.494) |
| Range | 12.667 - 64.667 | 11.000 - 54.000 | 7.333 - 54.667 | 8.000 - 49.333 |

**Figure S1.**
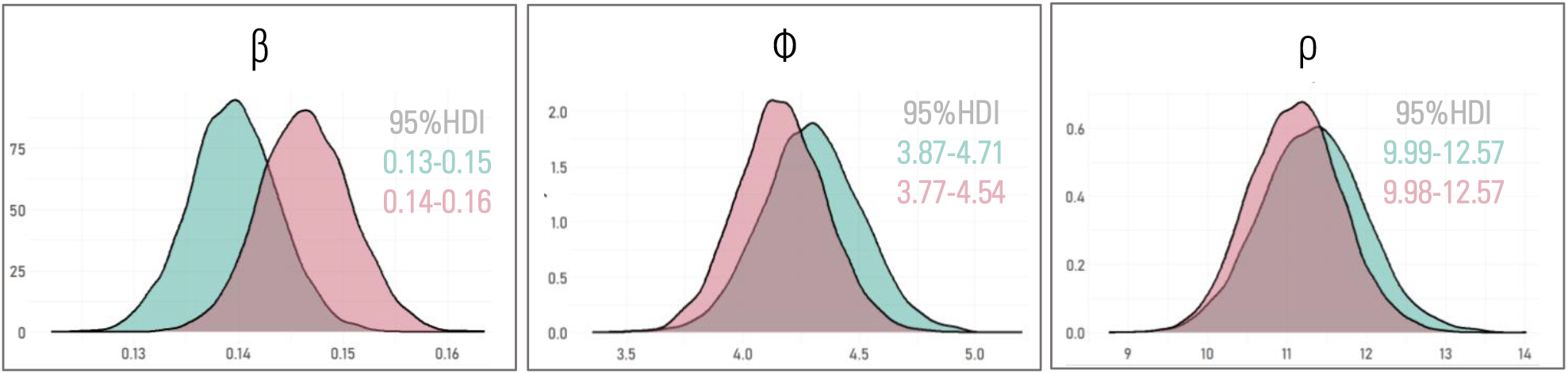
Posterior Distributions of Group-Level Mean Parameters derived from the winning model (BL + Bandit). Turquoise = PLC; Magenta = TYR; β = exploitation (value-based decision-making); Φ = directed exploration (via *bandit heuristic* exploration bonus); ρ = perseveration (stickiness; repetition choice at t-1). Numerical ranges in the plots show the 95% HDI for both conditions in their respective colors.

**Table S3.**
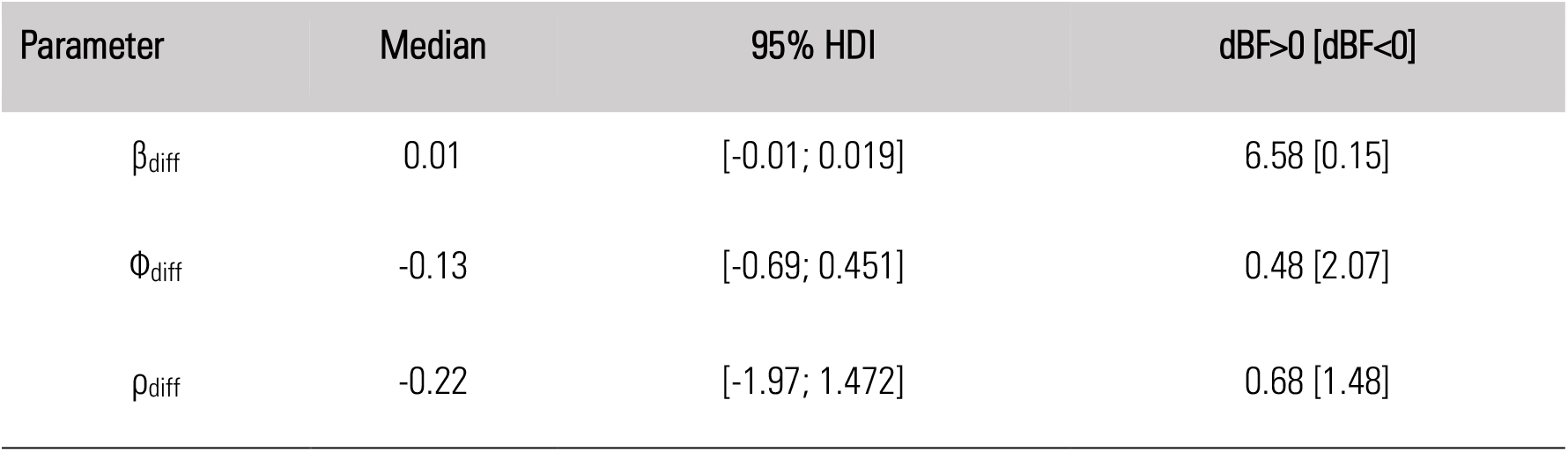
Summary of The Posterior Difference-Distribution. Parameters denote the group-level means’ difference distribution (TYR-PLC) of corresponding SoftMax parameters. dBF: directional Bayes Factor. β = exploitation (value-based decision-making); Φ = directed exploration (via *bandit heuristic* exploration bonus); ρ = perseveration (stickiness; repetition choice at t-1).

| Parameter | Median | 95% HDI | dBF>0 [dBF<0] |
| --- | --- | --- | --- |
| $\beta_{\text{diff}}$ | 0.01 | [-0.01; 0.019] | 6.58 [0.15] |
| $\Phi_{\text{diff}}$ | -0.13 | [-0.69; 0.451] | 0.48 [2.07] |
| $\rho_{\text{diff}}$ | -0.22 | [-1.97; 1.472] | 0.68 [1.48] |

**Figure S2.**
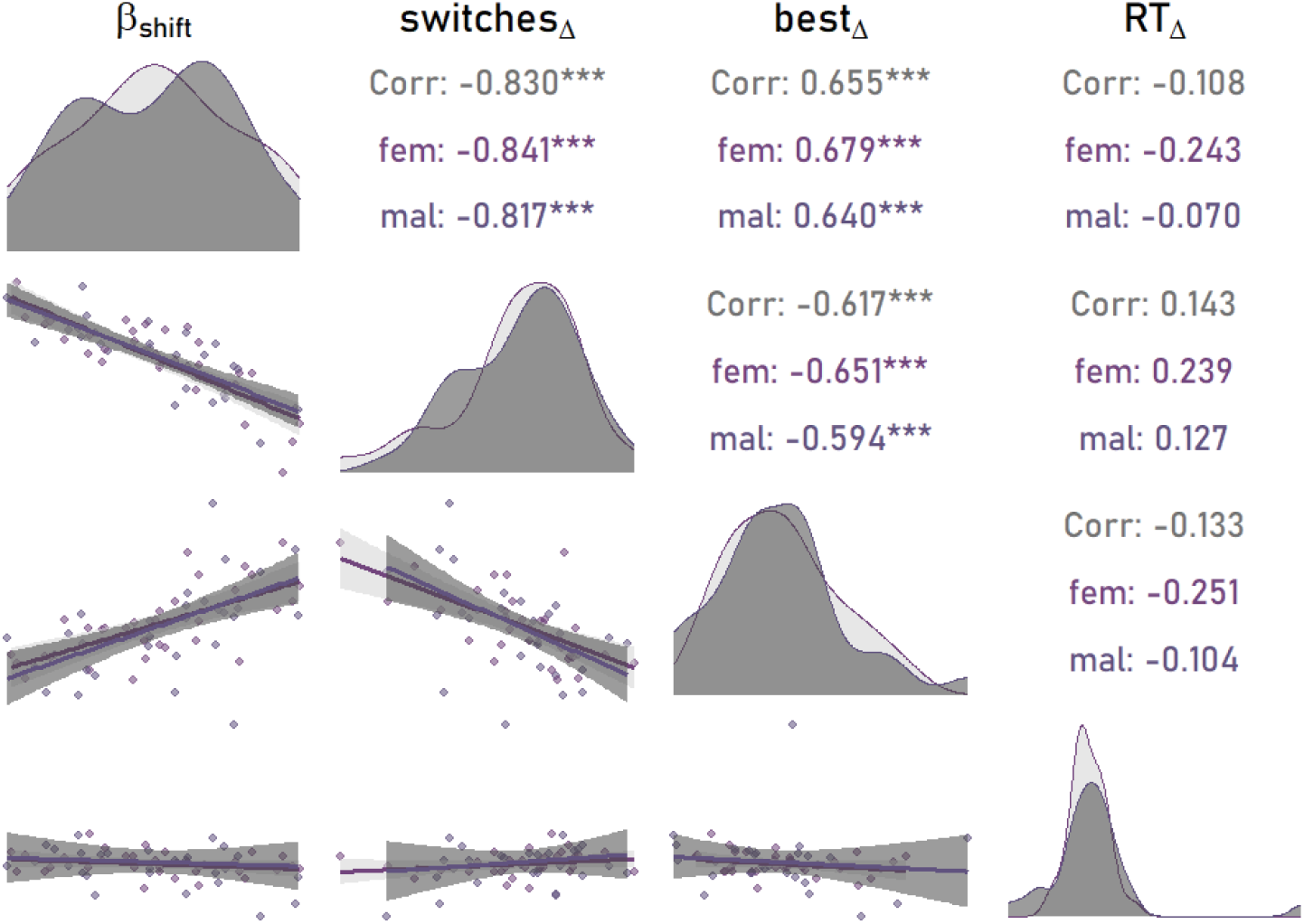
Association of Task Performance Measures with Supplementation-Effects on Value-Based Decision-Making. ß_shift_ : parameter depicting change in exploitation from PLC to TYR; switches_Δ_: difference in proportion of choice switches between conditions; best_Δ_: difference in proportion of highest-value choices; RT_Δ_: difference in median RTs between conditions. The diagonal shows the distribution of the associated parameter/measure separated by gender group. The lower triangular plots depict regression lines, surrounded by standard error intervals (shaded gray areas).

**Table S4.**
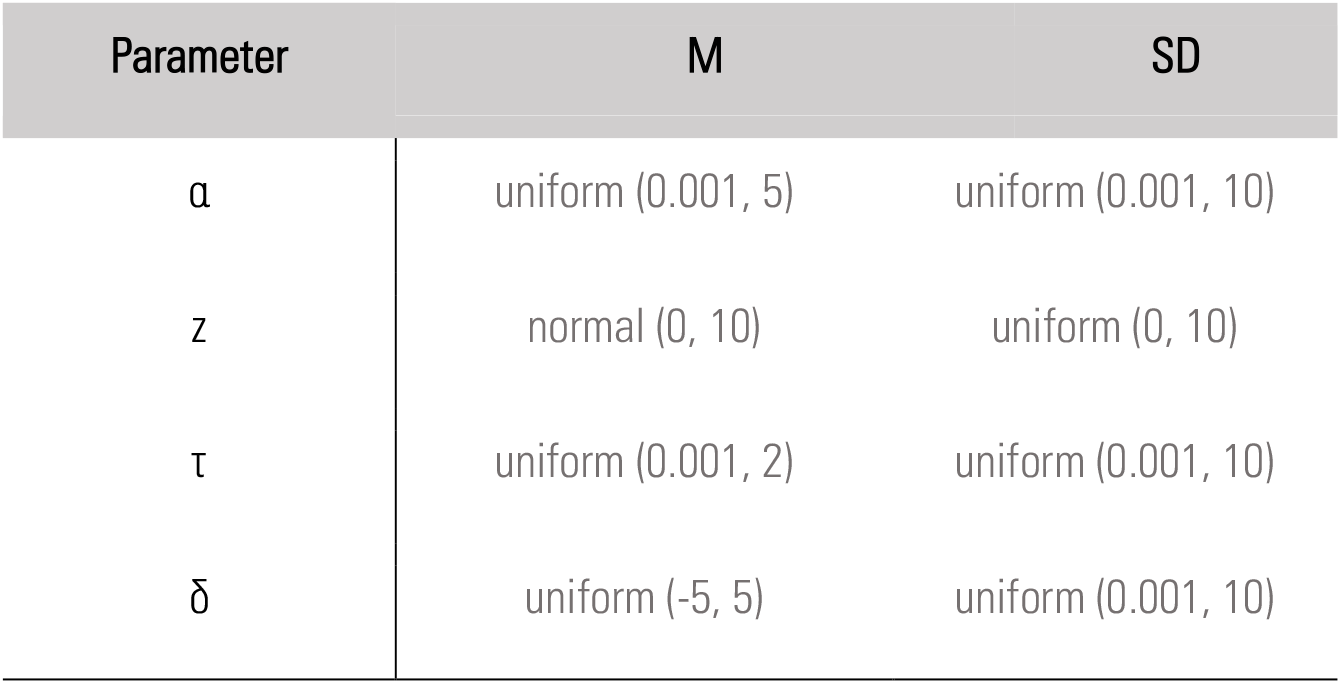
Priors for DDM Hyper-Parameters. For each subject-level parameter (first coloumn), M & SD denote priors for corresponding group-level parameters. Note that the bias parameter z was transformed to standard normal space within the model, so that priors listed below correspond to the untransformed parameter estimates.

| Parameter | M | SD |
| --- | --- | --- |
| $\alpha$ | uniform (0.001, 5) | uniform (0.001, 10) |
| $z$ | normal (0, 10) | uniform (0, 10) |
| $\tau$ | uniform (0.001, 2) | uniform (0.001, 10) |
| $\delta$ | uniform (-5, 5) | uniform (0.001, 10) |

**Figure S3.**
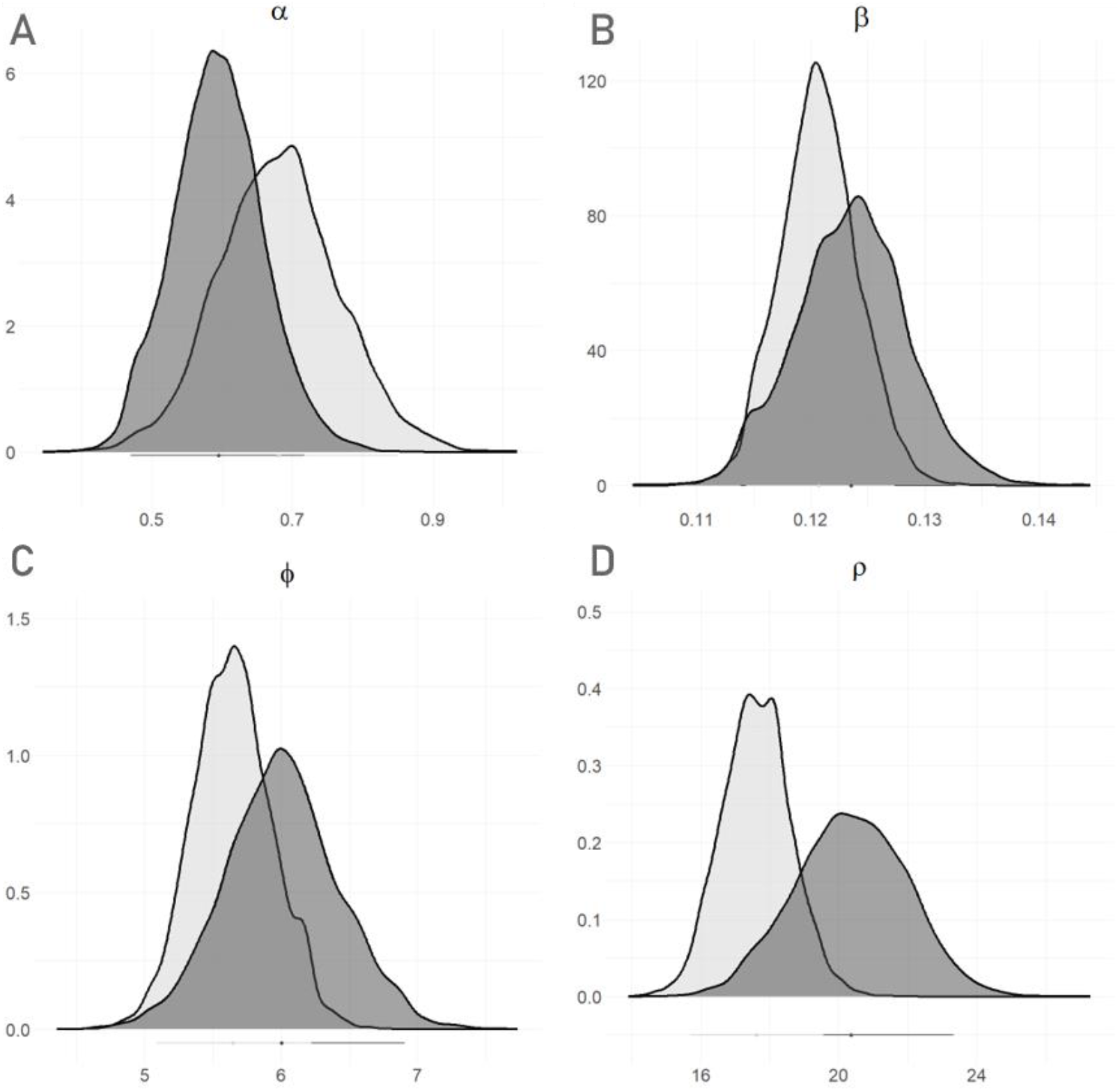
Posterior Distributions from the Exploratory Model Extension Separated by Gender Group. A) α: decay-rate parameter scaling the influence of past choice history; B) β: SoftMax parameter depicting exploitation; C) Φ: SoftMax parameter depicting directed exploration (via bandit-heuristic); D) ρ: SoftMax parameter depicting perseveration. Parameters were derived from the exploratory model extension of the winner model (BL + Bandit) by accounting for habitual perseveration (*higher order perseveration*). Light gray: female; Dark gray: male. Lines on the x-aches indicate the 95% HDI for each gender group in their respective colors.

**Table S5.**
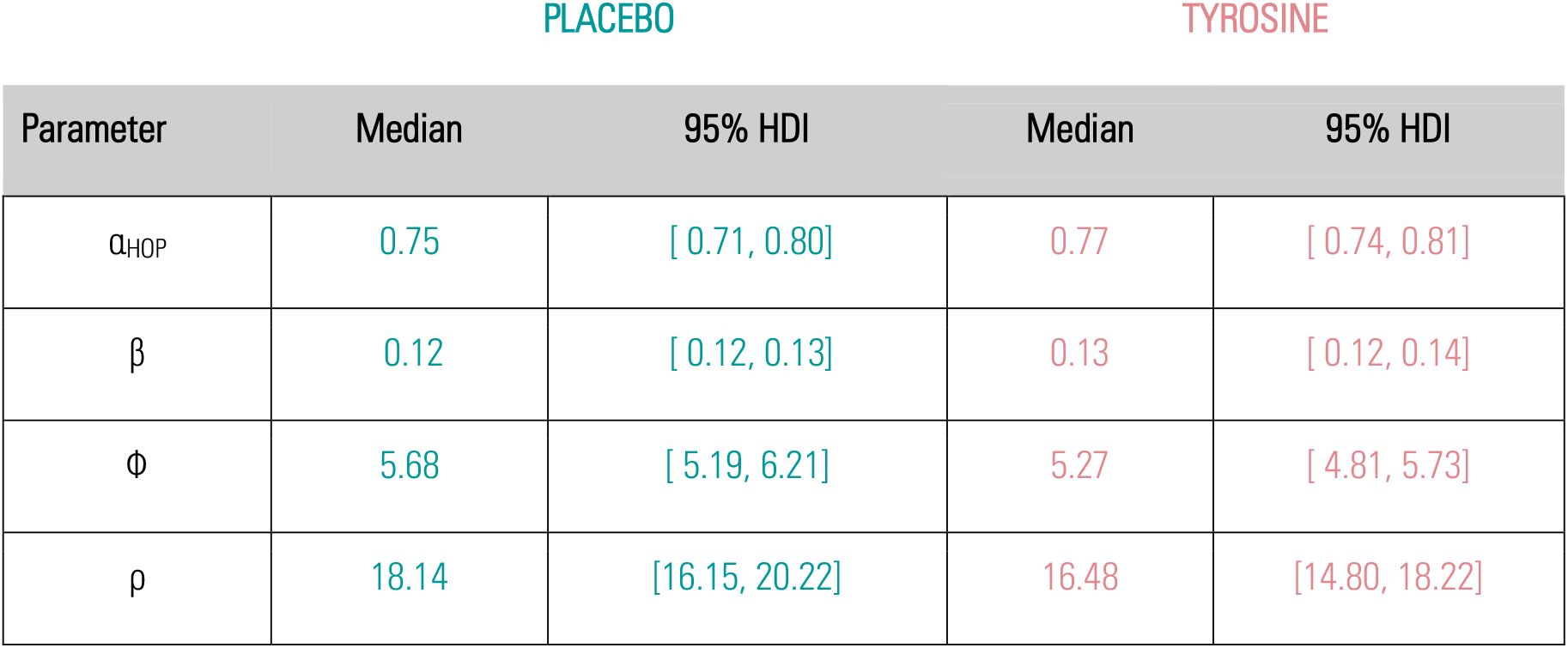
Posterior summary statistics of group level Mean parameter distributions from the exploratory HOP-model extension (BL + Bandit + Habit) separated by condition.

**Table S6.** Posterior Summary Statistics of TYR-effect parameters Based on Model BL + EXP + HABIT separated by gender group.

| Group | Parameter | Median | 95% HDI | dBF>0 [dBF<0] |
| --- | --- | --- | --- | --- |
| FEM | $\beta_{shift}$ | 0.13 | [-19.56, 19.54] | 1.02 [0.98] |
| | $\Phi_{shift}$ | -1.46 | [-19.14, 19.24] | 0.80 [1.24] |
| | $\rho_{shift}$ | -0.97 | [-18.75, 20.63] | 1.17 [0.85] |
| | $\alpha_{shift}$ | 0.07 | [-0.79, 0.29] | 1.05 [0.95] |
| MAL | $\beta_{shift}$ | 0.13 | [-19.57, 19.21] | 1.01 [0.98] |
| | $\Phi_{shift}$ | -0.85 | [-19.80, 19.95] | 0.86 [1.15] |
| | $\rho_{shift}$ | 0.15 | [-18.22, 19.18] | 1.00 [0.99] |
| | $\alpha_{shift}$ | -0.21 | [-0.76, 0.31] | 0.91 [1.10] |

**Figure S4.**
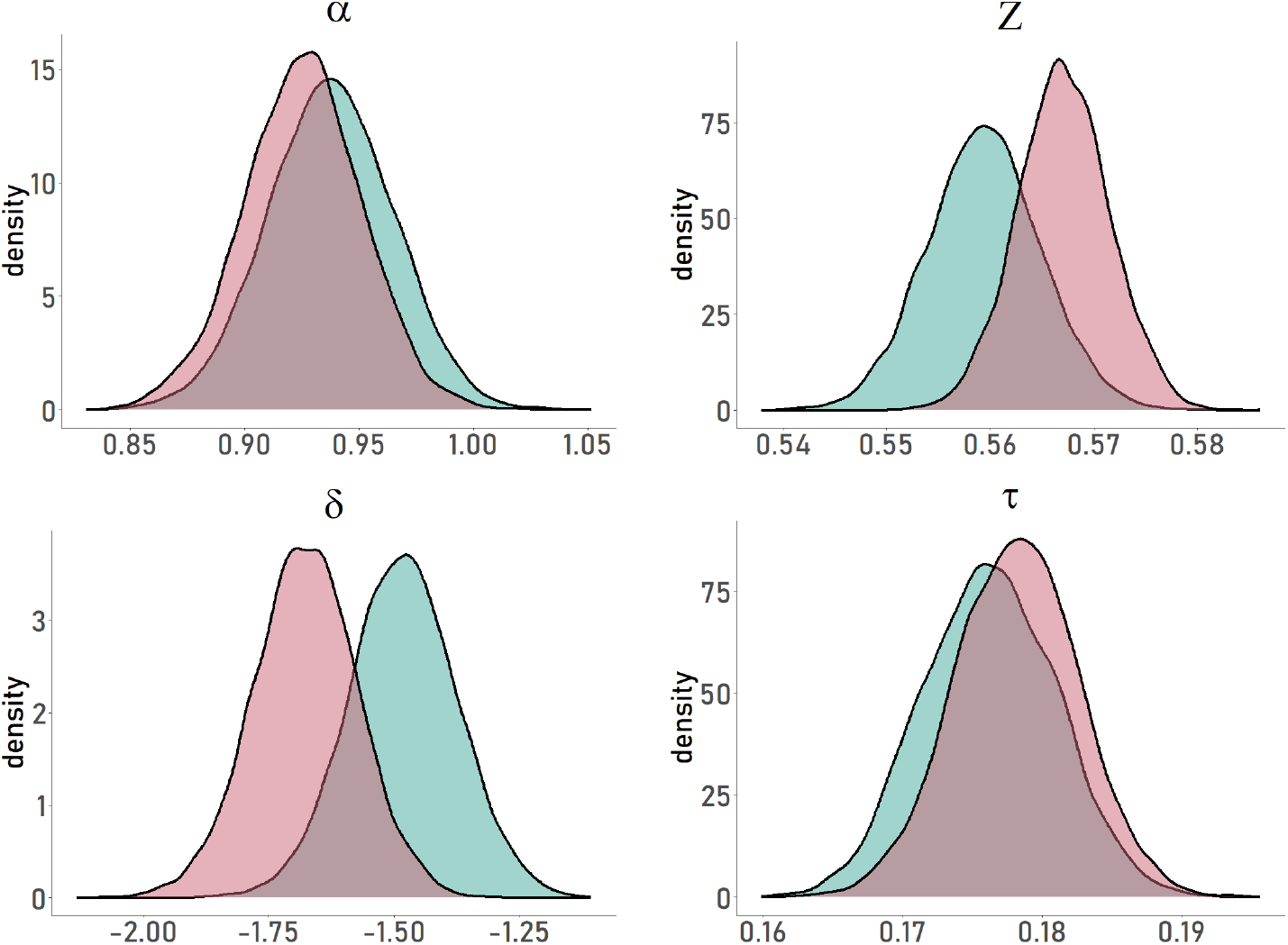
Posterior Distributions of DDM-Parameters Separated by Condition. Turquoise = PLC; magenta = TYR; α = boundary separation; z = bias; δ = drift rate; τ = non-decision time. Decision Boundaries were defined as 0 = stay (i.e. repetition of preceding choice); 1= switch (choice alteration).

**Table S7.** Single Condition Posterior Summary Statistics of DDM Fits.

| Parameter | PLACEBO |  | TYROSINE |  | DIFFERENCE |  |  |
| --- | --- | --- | --- | --- | --- | --- | --- |
|  | Median | 95% HDI | Median | 95% HDI | Median | 95% HDI | dBF>0 [dBF<0] |
| $\alpha$ | 0.82 | [ 0.79, 0.86] | 0.83 | [ 0.80, 0.87] | 0.010 | [-0.04, 0.06] | 1.58 [0.63] |
| $z$ | 0.56 | [ 0.55, 0.57] | 0.57 | [ 0.56, 0.58] | 0.008 | [-0.01, 0.02] | 6.21 [1.61] |
| $\tau$ | 0.18 | [ 0.17, 0.19] | 0.18 | [ 0.17, 0.19] | 0.002 | [-0.01, 0.01] | 1.58 [0.63] |
| $\delta$ | -1.48 | [-1.69, -1.26] | -1.68 | [-1.88, -1.47] | -0.190 | [-0.47, 0.12] | 0.11 [8.90] |

**Table S8.**
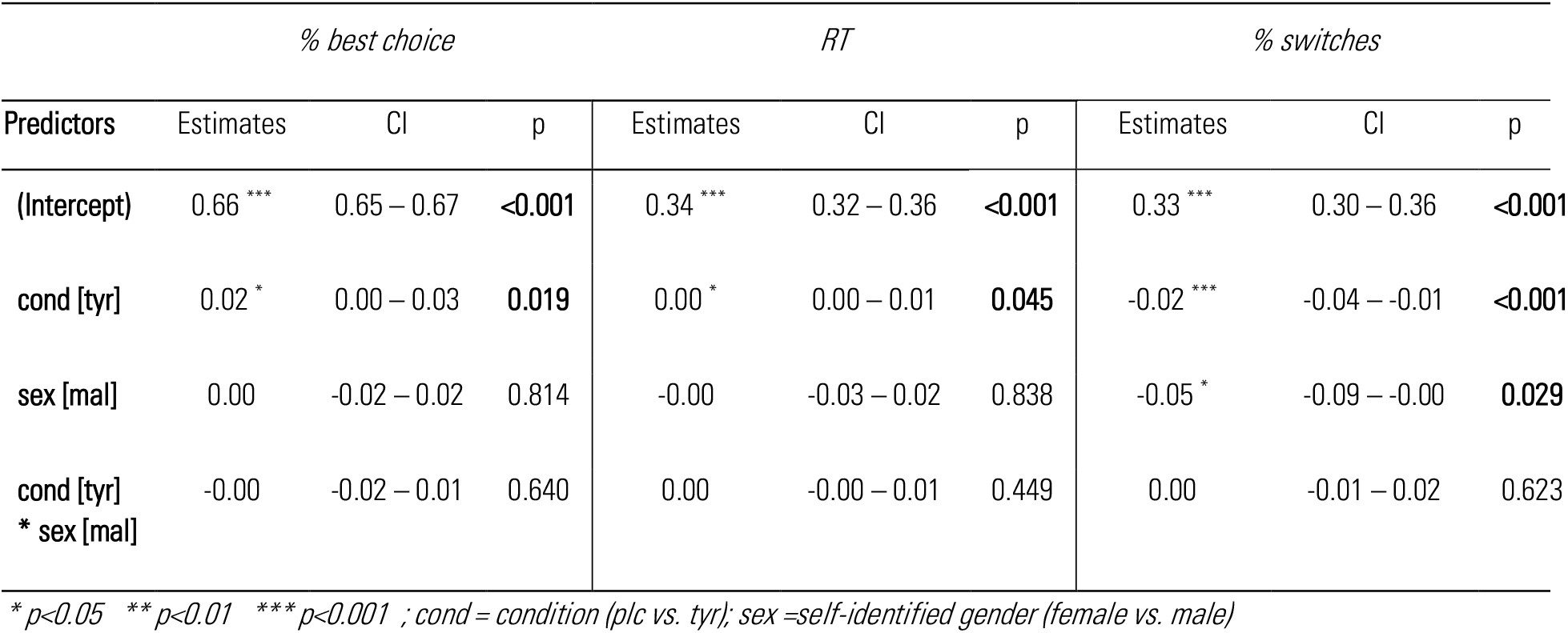
Control Regression Models Predicting Model-Agnostic Performance Indices While Accounting For Weight And Age.

